# EGF signalling in epithelial carcinoma cells utilizes preformed receptor homoclusters, with larger heteroclusters post activation

**DOI:** 10.1101/305292

**Authors:** Charlotte Fournier, Adam J. M. Wollman, Isabel Llorente-Garcia, Oliver Harriman, Djamila Ouarat, Jenny Wilding, Walter Bodmer, Mark C. Leake

## Abstract

Epidermal growth factor (EGF) signalling regulates cell growth, differentiation and proliferation in epithelium and EGF receptor (EGFR) overexpression has been reported in several carcinoma types. Structural and biochemical evidence suggests EGF binding stimulates EGFR monomer-dimer transitions, activating downstream signalling. However, mechanistic details of ligand binding to functional receptors in live cells remain contentious. We report real time single-molecule TIRF of human epithelial carcinoma cells with negligible native EGFR expression, transfected with GFP-tagged EGFR, before and after receptor activation with TMR-labelled EGF ligand. Fluorescently labelled EGFR and EGF are simultaneously tracked to 40nm precision to explore stoichiometry and spatiotemporal dynamics upon EGF binding. Using inhibitors that block binding to EGFR directly, or indirectly through HER2, our results indicate that pre-activated EGFR consists of preformed homoclusters, while larger heteroclusters including HER2 form upon activation. The relative stoichiometry of EGFR to EGF after binding peaks at 2, indicating negative cooperativity of EGFR activation.

---

The epidermal growth factor receptor (EGFR) is essential for normal growth and development of epithelial tissues and is a key component in several signaling pathways^1^. Aberrant signal transduction is a primary driver of many epithelial cancers, EGFR upregulation implicated in formation and progression of several carcinomas^2^. Human EGFR or ERBB1, (also denoted ‘ErB1’or ‘HER1’) is a 1,186 amino acid (aa) residue 170 kDa molecular weight protein^3^ belonging to a family of receptor tyrosine kinase (RTK) receptors with three additional members: ERBB2 (‘ErbB2’, ‘HER2’ or ‘neu’), ERBB3 (‘ErbB3’ or ΉER3’) and ERBB4 (‘ErbB4’ or ‘HER4’) expressed predominantly in the plasma membrane of epithelial cells^4^. EGFR has a 621aa extracellular region, divided into subdomains I-IV^5^. Domains I and III directly participate in ligand binding^6^, connected via a 23aa hydrophobic transmembrane α-helix to a 542aa cytoplasmic domain containing a 300aa tyrosine kinase^7^.

EGFR activation requires ligand binding, receptor-receptor interactions, and full activation of the tyrosine kinase^8^. At least 11 different ligands bind to the EGFR family, four to EGFR including EGF itself^9^. Prior to ligand binding the tyrosine kinase has low catalytic activity. Ligand binding results in full kinase activation through c-lobe interaction of an ‘activator’ and n-lobe ‘receiver’^10^. Subsequent autophosphorylation of intracellular tyrosine residues^11^ initiates intracellular reactions ultimately stimulating cellular growth, differentiation and proliferation^12^, terminated by internalization and proteolytic degradation of the receptor-ligand complex^13^.

The field has detailed insights concerning extracellular and intracellular interactions that contribute to signal transduction, however, there remains conflicting evidence concerning the *in vivo* composition of EGFR before and after activation and the role of higher order multimeric complexes of EGFR. Small angle X-ray scattering and isothermal titration calorimetry to EGFR’s isolated extracellular domain (sEGFR) suggests that EGF binds to an sEGFR monomer and that receptor dimerization involves subsequent association of two monomeric EGF-sEGFR^14^. Molecular weight determination by multi-angle laser light scattering suggests sEGFR is monomeric in solution but dimeric after addition of EGF^15^. Fluorescence anisotropy indicates a 1:1 binding ratio of EGF:sEGFR, with analytical ultracentrifugation suggesting the complex is comprised of 2(EGF-sEGFR)^16^. Structural evidence suggests activation is preceded by ligand binding to a receptor monomer^17–19^, and that EGF induces EGFR conformational change by removing interactions that auto-inhibit EGFR dimerization^20^. This model assumes that EGF binding increases the affinity for subsequent EGF to bind to the free EGFR subunit in the dimer (i.e. positively cooperative). However this is in conflict with EGF-EGFR binding studies of the full length receptor indicating that EGF binding reduces the affinity for subsequent EGF binding to the free EGFR subunit in the dimer ^21^ (i.e. negatively cooperative) mediated through the dimerization arm and intracellular juxta-membrane domain^22^. Recent structural studies of sEGFR in *Drosophila melanogaster* support a negatively cooperative model^23^, and it has been shown that EGFR dimers with a single bound EGF can be phosphorylated^24^. A predication from negative cooperativity is that EGFR:EGF bound complexes have a relative stoichiometry of 2:1 ^25^.

Chemical crosslinking and immunoprecipitation studies of full length receptors support a preformed dimer model^26^, suggesting that receptor dimerization is mechanistically decoupled from activation. Similarly, the first single-molecule fluorescence imaging studies on functional cell membranes suggested initial binding of one EGF molecule to a preformed EGFR dimer, rapidly followed by a second EGF to form a 2:2 complex^27^. Förster resonance energy transfer (FRET) studies subsequently reported preformed oligomeric EGFR^28^ supported by other live cell microscopy^29^, autocorrelation^30^, bimolecular fluorescence complementation (BiFC)^31^, fluorescence cross-correlation combined with FRET^32^, mobility measurements of quantum dot tagged EGFR^33^ and pixel brightness analysis of GFP-labeled EGFR^34^. Recent single-molecule photobleaching analysis suggests that EGFR forms oligomers prior to EGF binding^35^, and that EGFR clustering may be triggered at physiological EGF levels^36^, which contradicts live *Xenopous* oocyte studies that report a significant population of monomeric EGFR present before EGF activation^37^. The observed clustering of EGFR is not unique, but a general feature of cell membrane receptors in signal activation^38^. However, the EGFR clustering is nuanced in that it may involve cooperativity not only between monomer subunits of EGFR molecules in a dimer, i.e. an EGFR homodimer, but also between other ErbB receptor monomers of a different class, i.e. heterodimers^31,34^.

EGFR’s oligomeric state before and after activation under physiological conditions remains an open question due to technical limitations in obtaining simultaneous information for the relative stoichiometry of interacting receptors and ligands, the sensitive dependence of EGF expression levels on the EGFR state of oligomerization, the presence of both fluorescently labeled and natively unlabeled EGFR, and species-specific differences of model immortalized cell lines. Previous fluorescence microscopy studies on live cells have used non-epithelial immortalized rodent sources of mouse (BaF/3, B82, NIH/3T3) and hamster (CHO-K1). There have also been studies using human epidermoid carcinoma cells (A431, BT20, A549 and H460). All of these strains have measurable native levels of EGFR expression; in the case of the most commonly used A431 strain a staggering 2-6 million receptors per cell. Similarly, recent single-molecule investigations using transfected GFP-labeled EGFR in *Xenopus* oocytes may still exhibit appreciable expression levels of unlabeled native EGFR since their membrane surface forms microvilli in which EGF receptors localize^37^. Here, instead, we investigate a human epithelial carcinoma cell line, with no detectable native EGFR, to improve our understanding of EGF binding to EGFR in human cancer cells. We overcome previous technical limitations of simultaneous receptor and ligand measurements using single-molecule dual-colour total internal reflection fluorescence (TIRF) microscopy on live human colorectal carcinoma cells into which GFP-labelled EGFR has been stably transfected, coupled to real time nanoscale tracking of the red/orange dye tetramethylrhodamine (TMR) conjugated to EGF (Fig. 1a). We present results in the presence and absence of cetuximab^52^ or trastuzumab^41^, two popular immunotherapy antibodies which inhibit EGF signalling. We find that EGFR forms oligomeric clusters prior to EGF binding, with a mode peak stoichiometry of 6 EGFR molecules per cluster. After EGF binding, we observe clusters containing both EGFR and HER2. These are consistent with negative cooperativity for EGFR activation by EGF ^21^, resolving a key question in the field.

**Figure 1.**
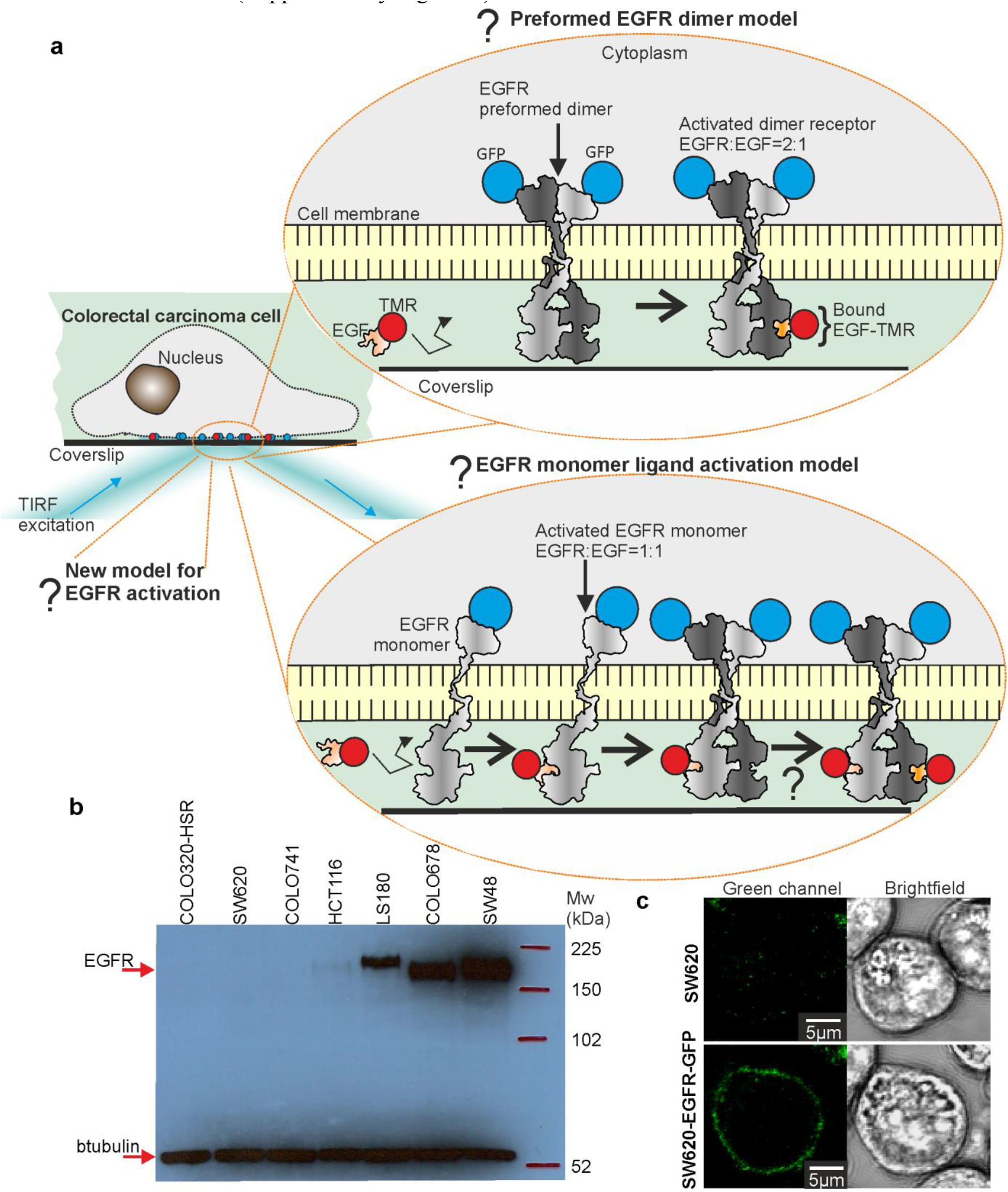
Visualizing functional EGF-EGFR complexes in human carcinoma cells. (**a**) Dual-colour TIRF applied to EGFR-GFP transfected human colorectal carcinoma cells with and without presence of fluorescently-labelled EGF-TMR. Several models to explain EGF activation of EGFR have been postulated, including ‘monomer’ and ‘preformed dimer’ models (EGFR structure PDB ID 1egf; EGFR monomer and dimer cartoons have been generated by manually combining separate structures with PDB ID values of 1nql, 1ivo, 2jwa, 1m17and 2gs6). (**b**) SDS-PAGE taken for several candidate colorectal carcinoma cell lines, indicating that SW620 COLO320-HSR (as opposed to COLO320-DM, its duplicate line) and COLO741 (later found to be a melanoma line and so not subsequently used here) have negligible native EGFR expression levels compared to positive controls of HCT116, LS180, COLO678 and SW48, shown to have intermediate EGFR expression levels. Note, there is a difference in apparent molecular weight for EGFR between LS180 and COLO678/SW48, most probably due to glycosylation. (**c**) Parental (non GFP) SW620 carcinoma cells show minimal autofluorescence in the green TIRF channel (left panel), while SW620-EGFR-GFP show membrane localization for EGFR-GFP (right panel).

## Results

### Construction of EGFR-GFP carcinoma cells

Human epithelial cell line SW620 was selected from an extensive colorectal carcinoma library for its undetectable EGFR expression as quantified by DNA microarray^42^ (Supplementary Fig. 1) and western blot (Fig. 1b). SW620 was stably transfected with plasmid pEGFR-EGFP-N1 to give SW620-EGFR-GFP (we denote EGFP throughout as simply ‘GFP’), GFP tagging the cytoplasmic domain far from the EGF binding site. Confocal microscopy of live cells confirmed membrane localization (Fig. 1c) with immunofluorescence on fixed cells demonstrating colocalization with EGFR (Supplementary Fig. 2a-d).

### TIRF optimized for single-molecule detection of EGF and EGFR

We optimized a bespoke dual-colour TIRF microscope (Supplementary Fig. 2e) for single-molecule detection using a fluorophore assay^43^ in which either GFP or EGF-TMR are conjugated to a glass coverslip using either IgG antibodies or derived Fab nanobody fragments with binding specificity to GFP or EGF (Supplementary Fig. 3a). We optimized imaging conditions to yield consistent fields of view containing fluorescent foci of GFP or EGF-TMR sampled at a video-rate of 30 ms per frame. Foci had a detectable brightness above background noise and a measured width (defined as half width at half maximum from their pixel intensity profile) in the range 250-300nm (in comparison to the measured point spread function (PSF) width of our microscope of 230nm). After ~1 s of continuous laser illumination foci exhibited irreversible step-wise photobleaching (Fig. 2a), indicative of single molecules of either GFP or EGF-TMR. Each focus had a brightness (summed pixel intensity integrated over each focus) of ~2,000 counts on our detector (Supplementary Fig. 3b). Although each IgG molecule contains two Fab sites, we saw no statistically significant difference in the number of two-step photobleach traces compared to Fab nanobody fragments, suggesting that GFP binding to an IgG Fab site may limit accessibility for a second GFP.

**Figure 2.**
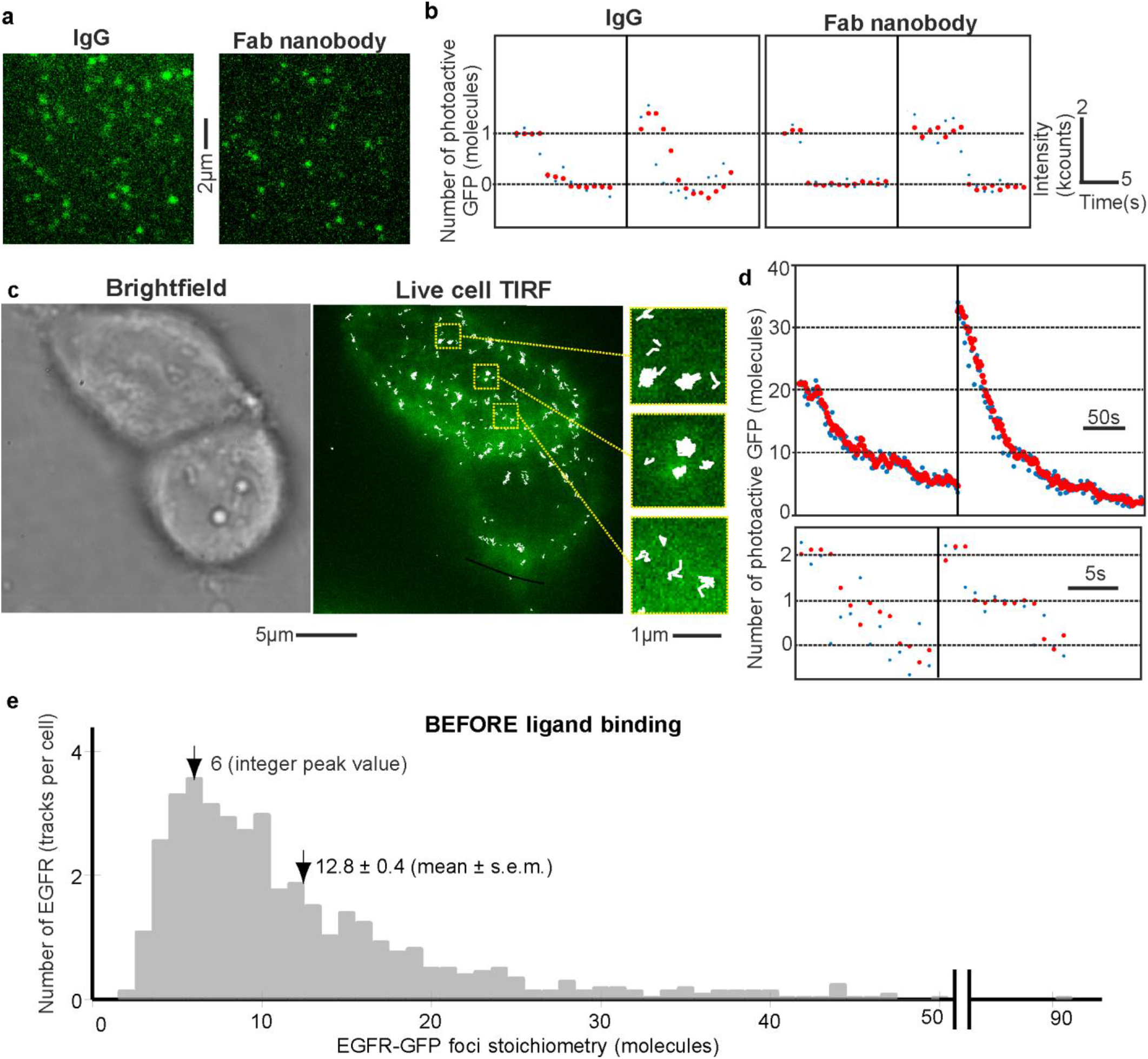
Stoichiometry of EGFR before EGF binding. (**a**) TIRF images of surface-immobilized GFP *in vitro* using IgG and Fab nanobody conjugation. (**b**) Example step-wise photobleach traces show raw (blue) and output data of an edge-preserving Chung-Kennedy filter^48,49^ (red), kcounts equivalent to counts on our camera detector x 10^3^. (**c**) Example of two nearby SW620-EGFR-GFP cells showing GFP fluorescence (green) and overlaid tracking output (white) with zoom-ins (inset). (**d**) Example photobleach traces from tracked EGFR-GFP foci which have stoichiometries of several tens of EGFR molecules (upper panel), down to an observed minimum of just two molecules (lower panel), raw and overlaid filtered data shown. (**e**) Distribution of EGFR-GFP foci stoichiometry before EGF activation, showing a modal peak at 6 and mean ~12.8 molecules. Data extracted from N=19 cells, detecting N=1,250 foci tracks, corresponding to mean of ~780 EGFR molecules per cell.

**Figure 3.**
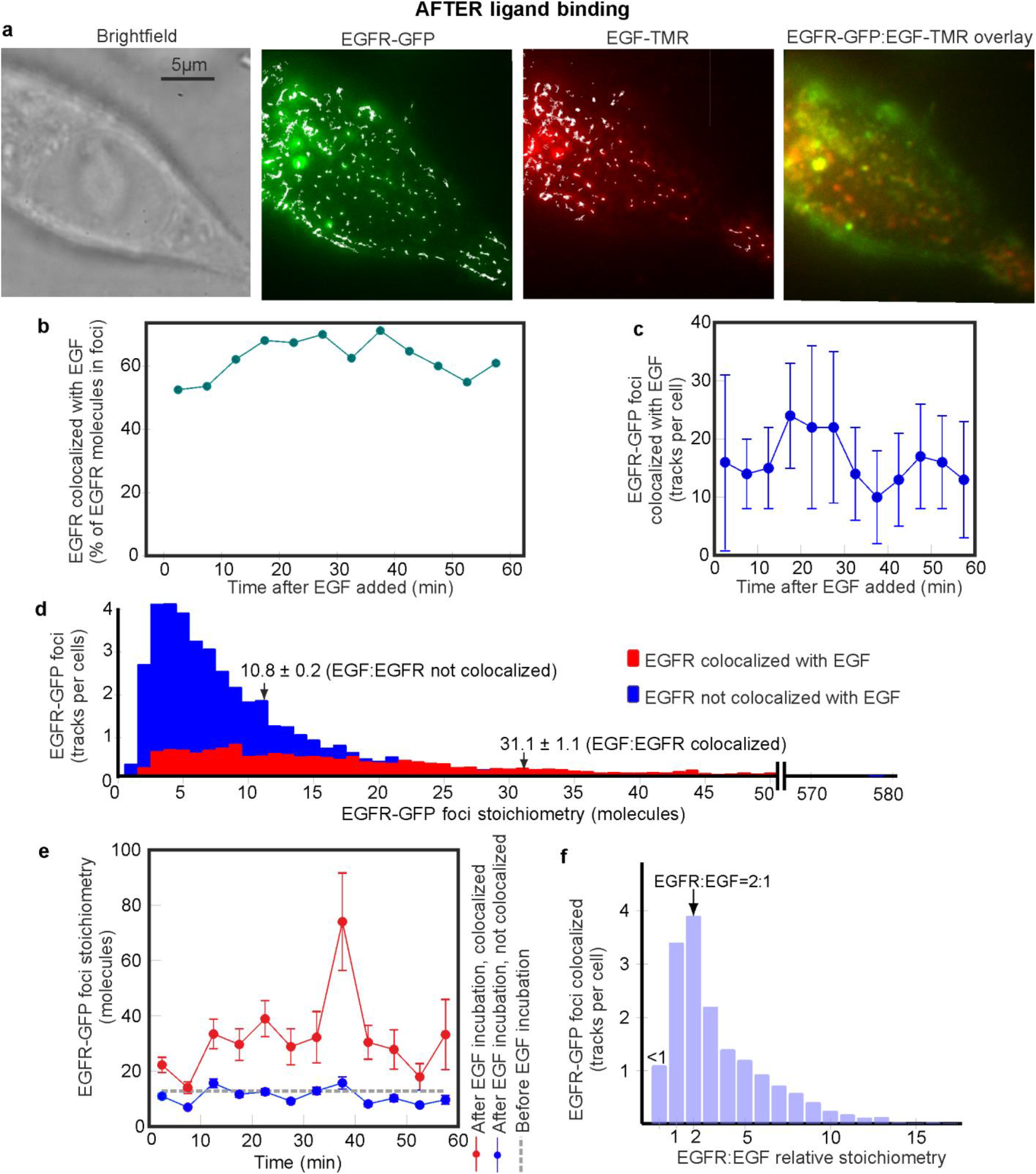
Effect of EGF binding on EGFR stoichiometry. (**a**) Brightfield and TIRF images of SW620-EGFR-GFP after adding EGF (~10 min incubation time point), GFP (green), TMR (red) and overlay images shown (yellow indicates high colocalization). (**b**) % of EGFR foci colocalized to EGF, (**c**) number of EGFR-EGF foci detected per cell (s.d. error bars). (**d**) EGFR-EGF foci stoichiometry (red) and isolated EGFR foci (blue) across all EGF incubation times, mean and s.e.m. indicated (arrows), and (**e**) as a function of incubation time (s.d. error bars). We categorized cells into 6 min interval bins resulting in N = 6-12 cells in each bin. (**f**) Distribution of relative stoichiometry of EGFR:EGF, integer bin widths, peak value at 2:1 indicated (arrow). Data extracted from a total of N = 119 cells.

### EGFR is oligomeric prior to EGF binding

To explore the architecture and dynamics of functional EGFR we used single-molecule TIRF on live SW620-EGFR-GFP cells. Prior to adding EGF in serum-free medium we observed several fluorescent foci in the GFP detection channel at a low surface density of 0.1-0.4 per µm^2^ in the plasma membrane (Fig. 2b and Supplementary Fig. 4). We tracked a mean of 66 ± 28 (s.d.) foci per cell and monitored their spatiotemporal dynamics over several seconds to a precision of ~40nm using bespoke software^44,45^, indicating a range of mobility (Supplementary Video 1). Foci widths were within ~10% of those observed for single GFP *in vitro*, however, brightness values were far greater. Foci brightness *vs*. time during tracking exhibited steps characteristic of stochastic photobleaching of one or more GFP within a single sampling time window (Fig. 2d), which we used to determine stoichiometry in terms of number of EGFR-GFP molecules present^43^. To estimate stoichiometry, initial foci brightness values were determined by interpolation to the start of each acquisition then divided by the *in vivo* brightness for a single GFP. To determine GFP brightness *in vivo* we quantified the mean foci brightness towards the end of each photobleach, when only one photoactive GFP molecule remained. Our analysis indicates that GFP brightness in a live cell is within 15% of that measured *in vitro* (Supplementary Fig. 2b). Previous live cell measurements using the same fluorescent protein indicate that the proportion of immature GFP is less than 15% of the total^56^. We measured a broad range of stoichiometry, both across different cells and within the same cell, of 2-90 EGFR molecules per fluorescent focus, with a peak integer value of 6 and associated mean of 12.8 ± 0.4 molecules (±s.e.m.) (Fig. 2e).

Since our microscope has the sensitivity to detect single GFP, one important conclusion is that there is no significant population of monomeric EGFR before adding EGF. The cell line has no detectable native EGFR expression, so our findings have consistency with a preformed dimer and/or oligomer model for EGFR^46^ as opposed to dimer formation being stimulated by EGF binding to monomeric EGFR, or where EGFR dimers are stabilized by two bound EGF^14^. We wondered if the observed stoichiometries could be due to random overlap of diffraction-limited images of individual EGFR-GFP foci. To address this question we modelled foci separation as a Poisson distribution^47^ (Methods), and used these to simulate apparent EGFR stoichiometries. We simulated monomeric, dimeric, and mixed oligomeric EGFR (monomers through to tetramers, suggested from a previous single-molecule live cell study^35^), all with poor agreement to the experimental data (Supplementary Fig. 5a, *R*^2^<0). We then tried a heuristic Monte Carlo overlap model (Methods) that simulated oligomeric EGFR whose stoichiometry was sampled randomly from a Poisson distribution with mean value equal to the peak of 6 that we observed, which resulted in a reasonable fit to the experimental distribution (Supplementary Fig. 5b, *R*^2^=0.4923).

### EGF binding to EGFR is negatively cooperative

To determine the effect of EGF binding on EGFR stoichiometry and spatiotemporal dynamics, live SW620-EGFR-GFP cells and non-GFP controls were kept in serum-free media for 12-24 h to minimize binding of any serum-based EGFR ligands. We visualized cells using dual-colour TIRF then added EGF-TMR, enabling simultaneous observation of EGFR and EGF in separate green and red colour channels respectively, before and after EGF activation. Excess EGF-TMR was retained in the sample chamber during imaging enabling observations over incubation times from 3-60 min. We observed a mean of 82 ± 36 EGFR foci tracks per cell across all incubation times, significantly higher than when EGF was absent. Colocalization of EGFR and EGF foci was determined using numerical integration between overlapping green and red channel foci^47^.

After EGF incubation from as little as a few minutes, colocalization between green and red channel foci was clearly detected (Fig. 3a, Supplementary Video 2 and Supplementary Fig. 6a). We estimated that 40 ± 18% of foci were colocalized EGFR-EGF when calculated across the full 60 min incubation, ~15 foci per cell or 64% of all EGFR molecules (Fig. 3b,c). EGFR-EGF foci had a statistically higher mean stoichiometry (Student’s *t*-test P<0.0001) of ~31 EGFR molecules compared to isolated receptors whose mean stoichiometry was ~11 EGFR molecules, consistent with measurements made before adding EGF indicating that effects from putative non-EGF ligands in the serum-free media were negligible (Fig. 3d, Table 1). The mean stoichiometry of isolated EGFR clusters remained roughly constant in the range ~8-14 molecules during incubation with EGF (Fig. 3e). The mean stoichiometry of EGFR-EGF clusters increased to ~32 EGFR molecules 10-15 min after adding EGF, up to a peak of ~70 EGFR molecules after ~40 min. At higher times EGFR endocytosis is prevalent^50^, consistent with observing some brighter EGFR foci in the main body of the cell, which may account for lower mean stoichiometry values of ~20-30 EGFR molecules per focus from ~40 min onwards.

**Table 1.** Mean EGFR foci stoichiometry values. Number of tracked foci in total (N foci) and individual cells (N cells) in datasets indicated. Biochemical interventions for added EGF (E), cetuximab (C), and trastuzumab (T) shown.

| Biochemical intervention |  |  | EGFR foci stoichiometry, uncolocalized |  | EGFR foci stoichiometry, colocalized with EGF |  | N cells |
| --- | --- | --- | --- | --- | --- | --- | --- |
| E | C | T | Mean ± s.e.m (molecules per EGFR focus) | N foci | Mean ± s.e.m (molecules per EGFR focus) | N foci | |
| - | - | - | 12.8±0.4 | 770 | X | X | 19 |
| + | - | - | 10.8±0.2 | 4,741 | 31.1±1.1 | 1,969 | 117 |
| - | + | - | 19.9±1.0 | 531 | X | X | 10 |
| - | - | + | 15.3±0.7 | 408 | X | X | 10 |
| + | + | - | 18.8±0.5 | 916 | 51.0±2.1 | 303 | 25 |
| + | - | + | 16.8±0.4 | 1,273 | 44.2±2.4 | 334 | 27 |

EGF-TMR quantified *in vitro* using conjugation to glass coverslips exhibited similar step-wise photobleaching as for GFP (Supplementary Fig. 3b). To determine the relative stoichiometry between EGFR and EGF when EGF is bound (i.e. the activated state) we measured red channel stoichiometry simultaneously to the green channel for EGF-EGFR foci. This analysis revealed a clear peak corresponding to a relative stoichiometry for EGFR:EGF of 2:1 (Fig. 3f, which pools data into integer width histogram bins). By using the measured variability in GFP and TMR brightness we estimate the error for the relative stoichiometry is ~0.7, in agreement with the half width at half maximum under the 2:1 peak, indicating that the apparent population in the 1:1 peak histogram bin is consistent with measurement error from the 2:1 population. Sub-dividing data by EGF incubation time revealed no significant shift in relative stoichiometry from the 2:1 peak (shown in kernel density estimations of Supplementary Fig. 6b where data has not been pooled into integer histogram bins). Before EGF-TMR was added in control experiments to the parental (non-GFP) strain we detected a small number of autofluorescent foci in red and green channels resulting in pseudo colocalization of ~2-3 tracks per cell (~3% of all colocalized foci). These pseudo colocalized tracks resulted in a small peak for the apparent relative stoichiometry in green:red colour channels equivalent to ~0.5:1 (Supplementary Fig. 6c), thus had a negligible impact on the measurements of the 2:1 peak. Adding EGF-TMR to this strain indicated foci detection levels in the red channel which were statistically indistinguishable (Student’s *t*-test P>0.05) to those measured in the absence of EGF-TMR (Fig. 4a).

**Figure 4.**
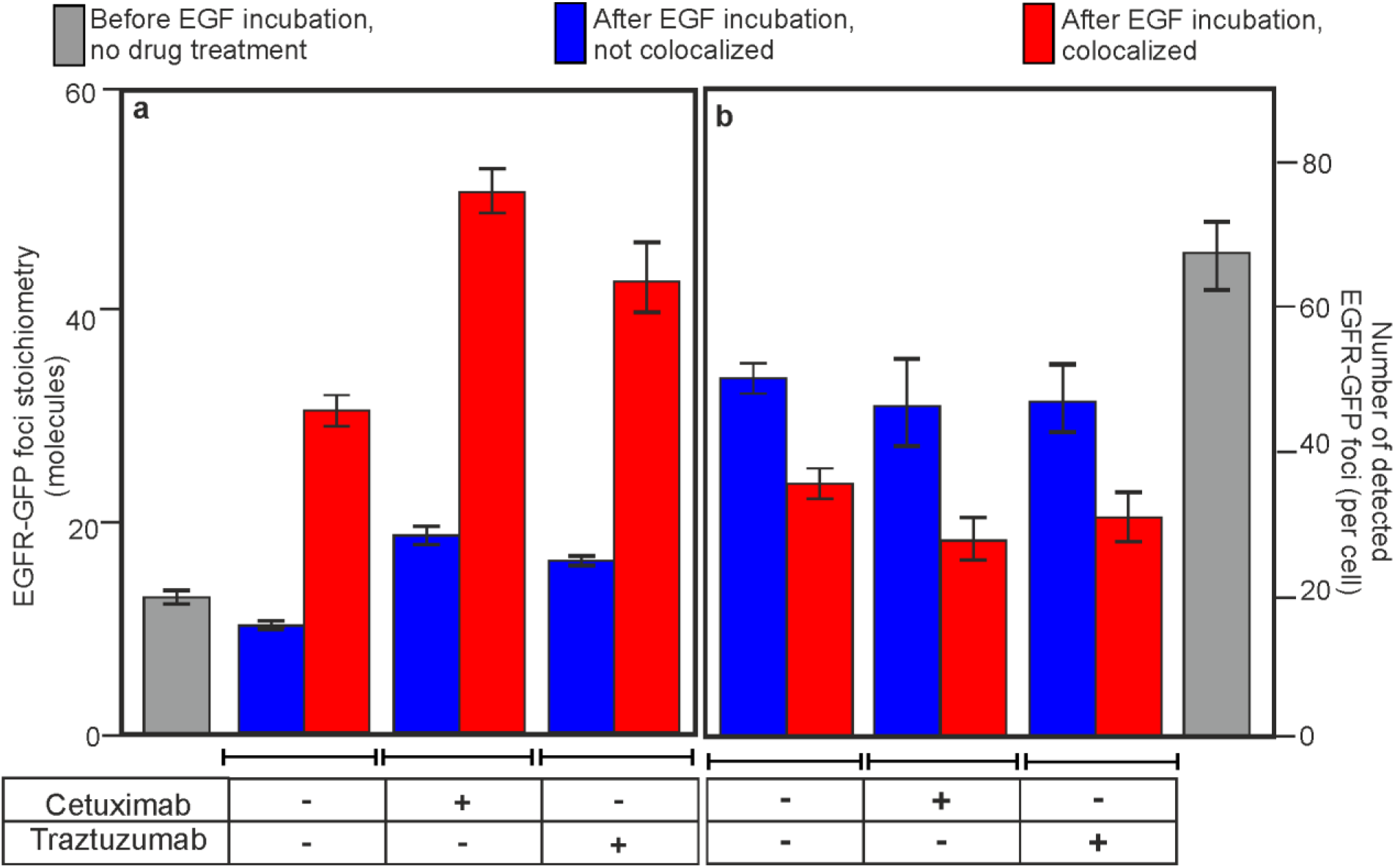
Effect of cetuximab and trastuzumab on EGF binding to EGFR. (**a**) Variation of mean EGFR-GFP foci stoichiometry, and (**b**) number of EGFR-GFP foci detected per cell. EGFR-EGF (red) and isolated EGFR foci (blue) are indicated for +/- addition of cetuximab and traztuzumab. Errror bars are s.d, number of cells per dataset in the range N =10 - 117.

Our findings indicate that the most likely receptor-ligand complex is a singly ligated EGFR dimer, consistent with a negatively cooperative mechanism for EGFR activation (Fig. 1a, upper schematic), i.e. a multiple EGF binding exclusion effect^21^. An alternative model consisting of initial EGF binding to monomeric EGFR to generate an activated state predisposed to form dimeric EGFR^17–19^ (Fig. 1a, lower schematic) predicts a significant 1:1 population, contrary to our observations. With this model, the proportion of 1:1 relative to 2:1 states might be expected to increase with longer EGF incubation times since there are putative steps directly dependent on the EGF on-rate, however, we observed no such dependence.

### EGFR clustering increases through direct and indirect EGF inhibition

To further understand the effect of EGF binding on EGFR clustering we performed live cell TIRF in the presence of cetuximab or trastuzumab. Cetuximab is a monoclonal antibody anti-cancer drug commonly used against neck and colorectal cancers in advanced stages to inhibit cell division and growth^51^, binding to domain III of the soluble extracellular region of EGFR, and believed to result in partial blockage of the EGF binding region as well inhibiting the receptor from adopting an extended conformation which may be required for EGFR dimerization^52^. Trastuzumab is also a monoclonal antibody anti-cancer drug, commonly used to treat breast cancer^52^, with similar effects of inhibiting cell division and growth, however, it does not bind directly to EGFR but instead to domain IV of the extracellular segment of HER2/neu, and its inhibitory action is believed to be related to the association of EGFR and HER2/neu in the plasma membrane^41^.

Before adding EGF we found that treatment with cetuximab or trastuzumab at concentration levels comparable to those used in cancer treatment resulted in a statistically significant increase in the mean stoichiometry of EGFR-GFP foci by ~25% and ~65% (Student’s *t*-test, P<0.0001) respectively (Fig. 4a), but with no effect on the number of detected EGFR-GFP foci per cell. Adding EGF resulted in ~20% fewer EGFR-EGF foci for cetuximab-or trastuzumab-treated cells compared to untreated cells (Fig. 4b).

The mean stoichiometry of EGFR-EGF foci in cetuximab and trastuzumab treatment datasets is 51 ± 2 and 44 ± 2 EGFR molecules per focus respectively, with the upper end having values of several hundred molecules (Fig. 5a, Table 1), consistent with previous qualitative observations that several different EGF pathway inhibitors increase EGRF clustering^53,54^. We also observed a shift to higher EGFR:EGF relative stoichiometry values for both cetuximab and trastuzumab treatments beyond the 2:1 peak observed for untreated cells (Fig. 5b).

**Figure 5.**
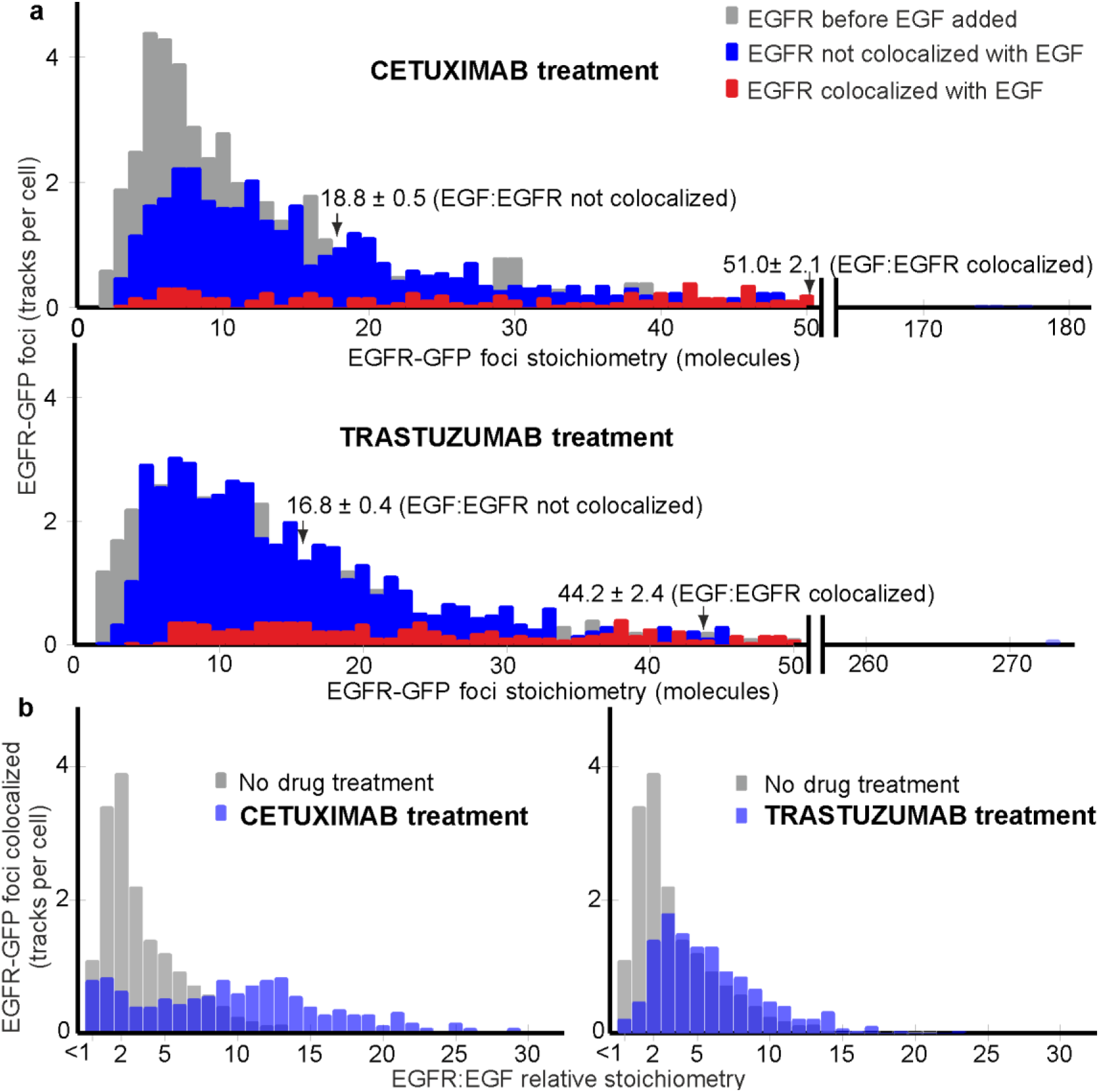
Effect of cetuximab or trastuzumab on EGFR foci stoichiometry. (**a**) Distribution of EGFR foci stoichiometry for cells treated with cetuximab or trastuzumab, showing pre (grey) and post EGF addition for EGFR-EGF (red) and isolated EGFR (blue) foci, data collated across 60 min EGF incubation time, mean and s.e.m. indicated (arrows). (**b**) EGFR:EGF relative stoichiometry of EGFR-EGF foci for drug-treated cells (blue) contrasted against no drug treatment (grey). Number of cells per dataset in the range N =10 - 117.

### EGF can trigger formation of larger EGFR heteroclusters

Tracking of EGFR foci indicated complex mobility in the plasma membrane: Brownian diffusion up to tracking time intervals of ~100 ms (Fig. 6a), transiently confined diffusion into zones of diameter ~400-500nm at time intervals of ~100-600 ms, and Brownian diffusion for time intervals >600 ms (shown indicatively in Supplementary Fig. 7a for the average mean square displacement up to time intervals of several seconds) similar to complicated patterns of diffusion observed previously for membrane proteins interacting with the cytoskeleton^55^. Using the initial gradient of the mean square displacement with respect to tracking time interval for each track we determined the apparent microscopic diffusion coefficient and correlated this against EGFR foci stoichiometry. We used a simple model based on the Stokes-Einstein relation, that the cross-sectional area of an EGFR cluster parallel to the plasma membrane scales linearly with the number of EGFR dimers present. The model assumes that the diffusion coefficient *D* is given by *k_B_T/γ* where *k_B_* is Boltzmann’s constant, *T* the absolute temperature and *γ* the frictional drag of the whole EGFR cluster in the membrane. The frictional drag is proportional to the effective radius of the EGFR cluster, which implies that *D* is proportional to the reciprocal of the square root of the stoichiometry. Our model results in reasonable agreement for data corresponding to pre and post EGF incubation (Fig. 6b).

**Figure 6.**
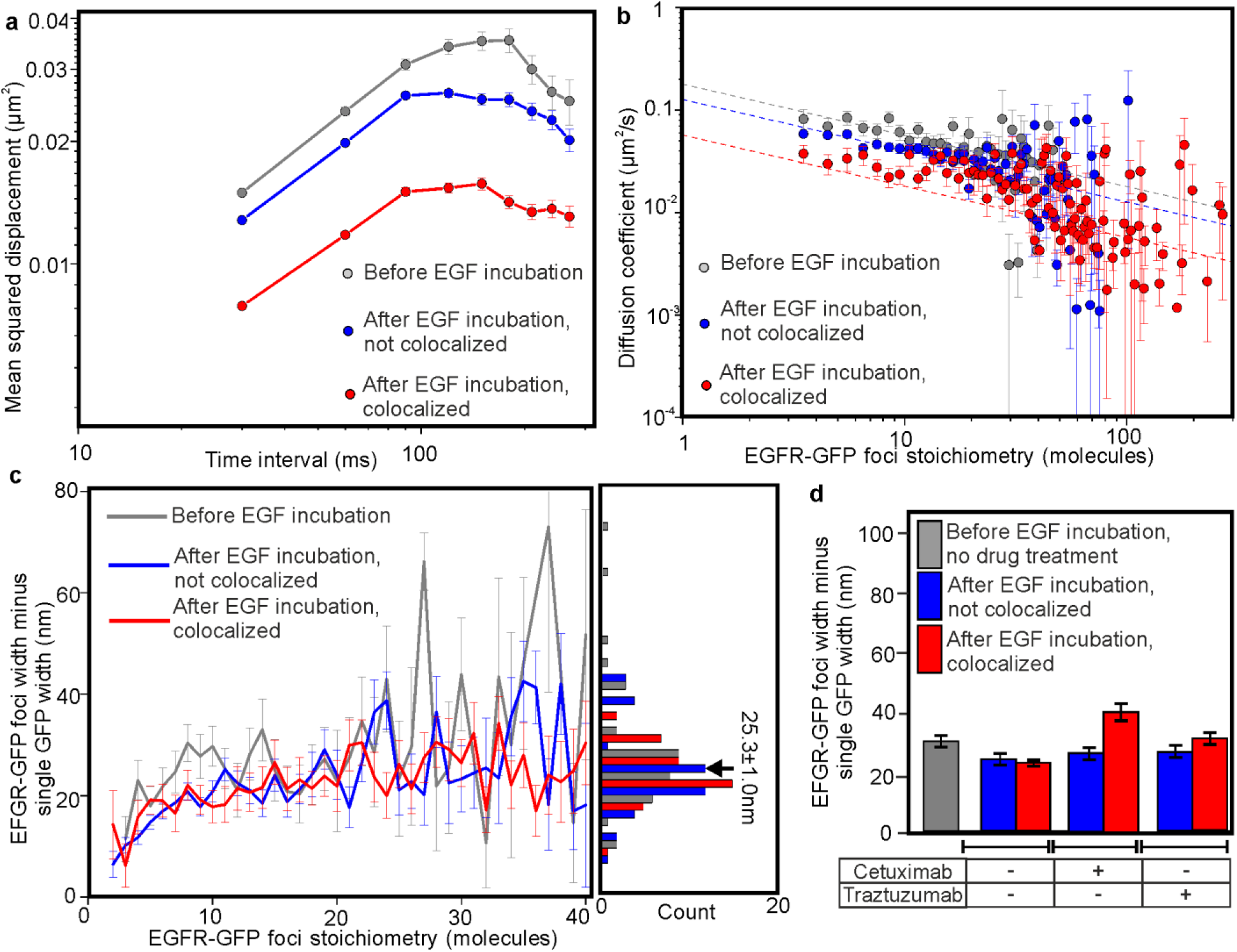
EGFR foci mobility depends on stoichiometry and EGF binding. (**a**) Log-log plot for average mean squared displacement for time intervals of 300 ms or less, and (**b**) log-log plot for apparent microscopic diffusion coefficient *D* with EGFR stoichiometry S, fits shown to Stokes-Einstein model assuming *D*~*S*^-1/2^ (dashed lines). (**c**) EGFR-GFP foci width minus the width of a single GFP *vs*. stoichiometry, and associated histogram, mean and s.e.m. for all datasets combined indicated (arrow). PreEGF incubation (grey, from N=770 foci, taken from N=19 cells) and post EGF incubation for EGFR-EGF (red, from N=1,969 foci, taken from number N=117 cells) and isolated EGFR (blue, from N=1,741 foci, taken from N=117 cells) foci shown, s.e.m. error bars. (**d**) Histograms EGFR-GFP mean foci width minus width of a single GFP. Pre EGF incubation for cells untreated with drugs (grey, from N=1,252 foci, taken from N=19 cells); cetuximab-treated cells post EGF incubation for EGFR-EGF (red, from N=151 foci, taken from N=10 cells) and isolated EGFR (blue, from N=1,253 foci, taken from N=10 cells) foci shown; trastuzumab-treated cells post EGF incubation for EGFR-EGF (red, from N=263 foci, taken from N=27 cells) and isolated EGFR (blue, from N=1,479 foci, taken from N=27 cells) foci shown; s.e.m. error bars.

We quantified EGFR-GFP foci widths by performing intensity profile analysis on background-corrected pixel values over each foci image^56^, and compared this with measurements obtained from single GFP *in vitro*, as a function of foci stoichiometry *S* (Fig. 6c). In all cases the mean EGFR-GFP foci width was greater than that of single GFP, which increased with the number of EGFR-GFP molecules present, consistent with a spatially extended structure. The dependence of this increase could be modelled with a heuristic power law relation *S^a^* with optimized exponent *a* of 0.27 ± 0.04 (s.e.m.) showing no dependence with EGF activation (Supplementary Fig. 7b), with a mean for all pooled data of 25.3 ± 1.0nm (s.e.m.). At the low end of *S* the increase in foci width minus single GFP width was ~11-12nm, while at the high end, corresponding in some cases to several hundred EGFR per focus, the increase in width was 30-40nm. Foci widths indicated no significant differences upon addition of either cetuximab or trastuzumab prior to addition of EGF (P>0.05), however, we observed an increase of ~50% for EGFR-EGF foci for cetuximab-treated cells (P<0.001) (Fig. 6d). Cells treated with cetuximab or trastuzumab exhibited a similar shape for the mean square displacement *vs*. time interval relation to untreated cells (Fig. 7a). Both treatment groups also showed reasonable agreement to a Stokes-Einstein model for diffusion, for before and after addition of EGF (Fig. 7b).

**Figure 7.**
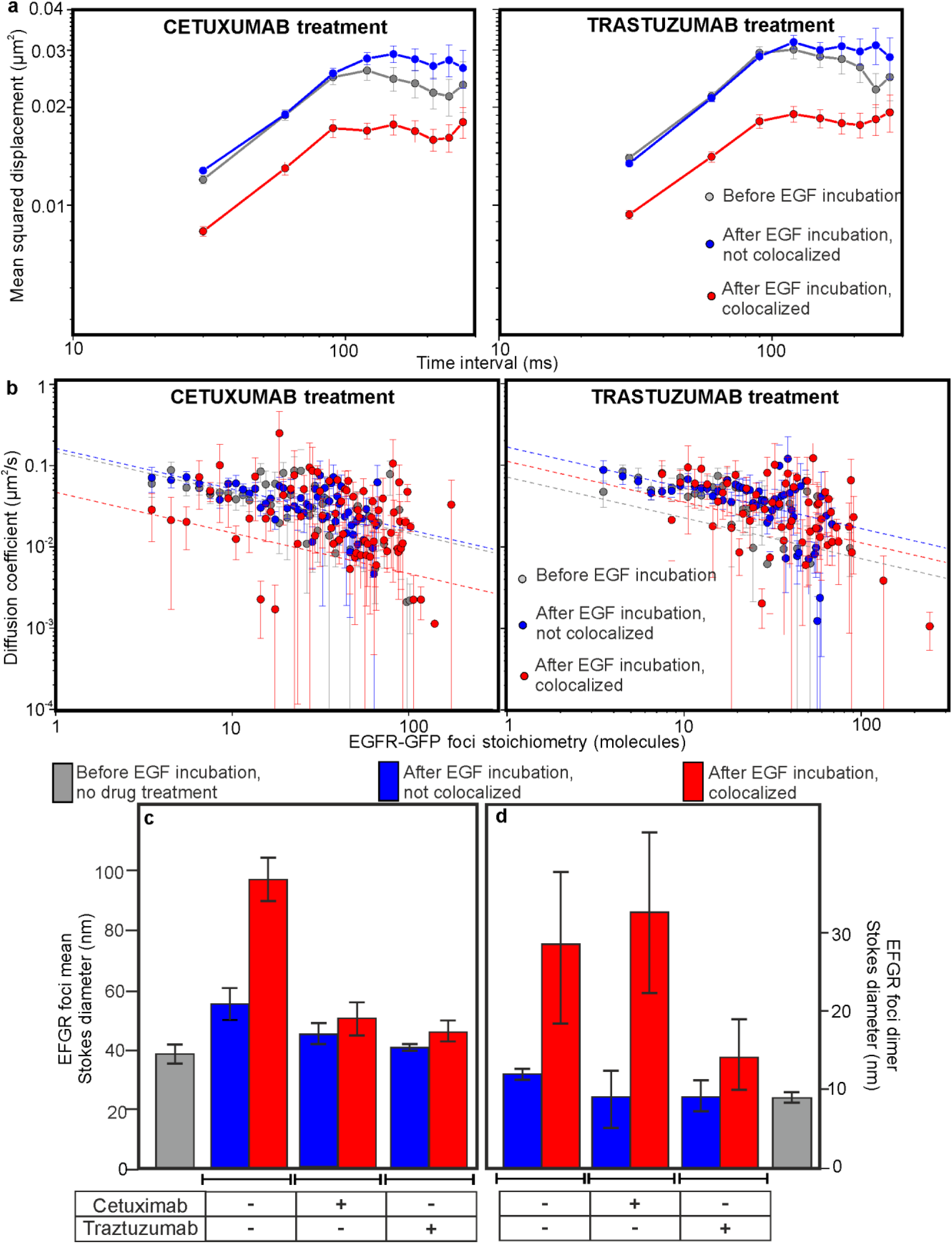
EGFR mobility can be affected by EGF inhibitors. (**a**) Log-log plots for average mean squared displacement for time intervals of 300 ms or less, and (**b**) log-log plots for variation of apparent microscopic diffusion coefficient *D* with EGFR stoichiometry S, fits shown to Stokes-Einstein model assuming *D*~*S*^-1/2^ (dashed lines) for cetuximab-and trastuzumab-treated cells. (**c**) Histogram of mean Stokes diameter, and (**d**) equivalent diameter values extrapolated for EGFR dimeric foci using same datasets as for Fig. 6d, s.e.m. error bars.

We used *D* to directly estimate the physical diameter of EGFR foci. A full analytical treatment models diffusion of membrane protein complexes as cylinders with their long axis perpendicular to the membrane surface^57^ requiring precise knowledge of local membrane thickness, however, here we simplified the analysis by calculating the diameter of the equivalent Stokes sphere to generate indicative values of drag length scale. We approximated the frictional drag by *3πηd* where *d* is the sphere diameter, assuming that drag contributions from the extracellular and cytoplasmic components are negligible since the kinematic viscosity *η* in the plasma membrane is higher by 2-3 orders of magnitude^58^. Using a consensus value of ~270 cP for the effective plasma membrane viscosity, estimated from human cell lines using high precision nanoscale viscosity probes^59^, indicates a mean Stokes diameter of ~40-60nm for isolated EGFR. EGFR-EGF foci had a mean Stokes diameter of closer to ~90nm, reduced back to the level for isolated EGFR to within experimental error upon treatment of cetuximab or trastuzumab (Fig. 7c).

We then used Stokes-Einstein fits to determine the Stokes diameter corresponding to the EGFR dimer (i.e. a stoichiometry of precisely 2), which indicated values in the range ~7-10nm for isolated EGFR across the cetuximab, trastuzumab and untreated cell datasets (Fig. 7d), broadly consistent with expectations from the crystal structure of dimeric EGFR^17,18^. EGF-EGFR foci corresponding to an EGFR stoichiometry of 2 had Stokes diameters of ~30nm, which were unaffected by cetuximab but reduced by a factor of ~2 almost to the level of isolated EGFR dimers by trastuzumab (Fig. 7d).

The Stokes diameter for an EGFR cluster is a measure of visible EGFR-GFP content plus any unlabelled associated protein contributing to overall frictional drag. Here, the proportion of non-fluorescent EGFR is low. However, other studies have suggested that EGFR forms heterocomplexes with other RTK receptors^31–33,60^. Here, we observe that treatment with the HER2-binder trastuzumab results in a similar measured Stokes diameter for EGF-EGFR foci to that for isolated EGFR dimers, suggesting that HER2 may form heterocomplexes with EGFR following EGF binding (Fig. 8, left panel). Also, since the mean Stokes diameter of EGFR-EFR foci of ~90nm corresponds to a stoichiometry ~32 EGFR molecules, i.e. 16 EGFR dimers, then the average diameter associated with a single EGFR dimer which can account for the same cluster area is ~22nm, greater than the measured diameter of an EGFR dimer from crystal structures^17,18^ by a factor of ~2. In other words, the observed Stokes diameter could be explained if EGFR-GFP dimers associate in a 1:1 relative stoichiometry with unlabelled HER2 dimers of similar same size and structure.

**Figure 8.**
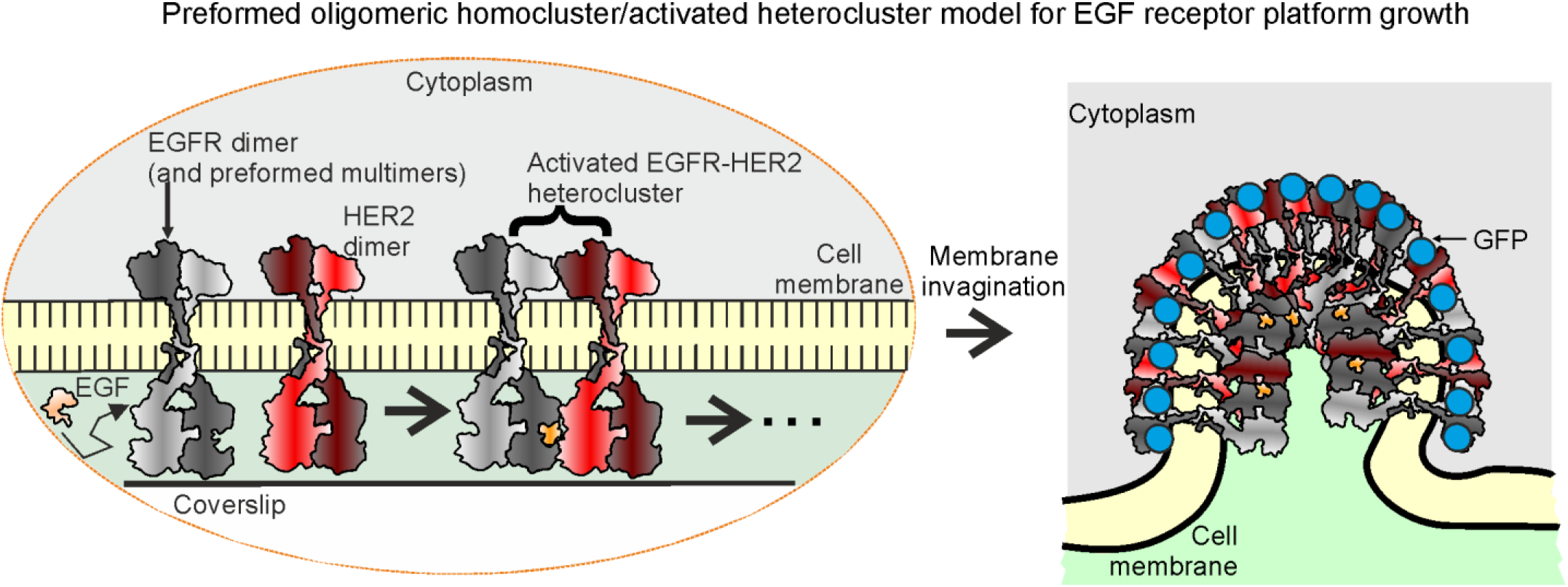
Activated EGFR nanoclusters grow in platforms containing heteroclusters of EGFR and HER2. Schematic illustrating how HER2 and EGFR dimers may be associated following EGFR activation by EGF (left panel) and how further oligomerization may result in local membrane invagination to form hetero receptor ‘platforms’ of several tens ofnm diameter.

An additional phenomenon to consider is plasma membrane invagination as EGFR clusters grow, ultimately culminating in a clathrin-coated vesicle inside the cytoplasm. Since the visible focus that we detect in TIRF corresponds to the GFP localization pattern in the invaginated membrane projected laterally onto our camera detector then its apparent visible diameter might appear to approach an asymptotic plateau with respect to EGFR-GFP stoichiometry (Fig. 8, right panel), which is broadly what we observed (Fig. 6c).

## Discussion

In summary, data acquired using genetics, cell biology, biochemical and biophysical techniques, in particular single-molecule imaging with tracking to 40nm precision and quantitative molecular and mobility analysis on live bowel carcinoma cells, suggest that preformed homo-oligomeric EGFR is present in the plasma membrane prior to EGF activation, comprising predominantly clusters of EGFR dimers. These observations are consistent with negative cooperativity of EGF binding to EGFR. Using a GFP probe on EGFR in combination with the spectrally distinct TMR tagged to EGF enabled unparalleled insight into the molecular stoichiometry, mobility and kinetics of single functional EGFR clusters in their pre and post activated states. Our observations indicate that the most prevalent EGFR complex in the absence of bound EGF is a hexamer, though with higher order oligomers also present extending to clusters containing up to ~90 molecules. We find that activation by EGF results in a shift to higher cluster stoichiometry, contrary to earlier speculation from experiments in which just EGF was fluorescently labelled suggesting tetrameric EGFR is the most likely multimeric state^36^. We observe that the action of cetuximab and trastuzumab, commonly used anti-cancer drugs, results in increases in the mean EGFR content of receptor clusters by a factor of 3-5. In addition, our findings suggest that EGF activation generates hetero clusters of EGFR and HER2, a response which results in the formation of super-clusters whose effective diameter is up to ~90nm.

Our findings clearly indicate that EGFR is clustered both before and after EGF activation, consistent with observations from earlier AFM imaging experiments which probed the surface morphology of the human lung adenocarcinoma cell line A549, known to have high levels of EGFR expression in the cell membrane^61^. This study suggested that half of the EGFR clusters quantified had a diameter in the range 20-70nm in the pre-activated state, and 35-105nm post activation, comparable with our light microscopy measurements. However, we find important differences with respect to recent single-molecule studies of EGFR activation^27,34–37^. We observe no significant monomeric population of EGFR before or after EGFR activation, despite having the sensitivity to detect single GFP under our imaging conditions, though we do observe the presence of single EGF-TMR molecules associated with multimeric EGFR clusters. Two key improvements in our study are that: (i) we specifically selected a human carcinoma cell strain with negligible native EGFR expression, whereas earlier single-molecule studies utilised cell lines likely to have much higher EGFR expression; (ii) unlike earlier studies we have definitive spatial information concerning the localization of EGFR and EGF simultaneously and so have a high level of confidence concerning the effects of EGF binding on the stoichiometry of specific individual EGFR foci. In single-molecule experiments in which EGFR is labelled with a fluorescent protein reporter for which there is some expression of native EGFR even if low^34,37^ then apparently monomeric EGFR foci may inevitably be detected even if a functional cluster has a higher stoichiometry, due to mixing of unlabelled and labelled EGFR molecules. In single-molecule experiments in which labelled EGF is not imaged simultaneously with labelled EGFR^27,35,36^ then no direct inference can be made as to the relative stoichiometry of associated clusters.

The lack of evidence in our experiments for a monomeric EGFR population coupled to a distinct peak of 2:1 for the EGFR:EGF relative stoichiometry as determined on a unique cluster-by-cluster basis provides clear evidence in support of negatively cooperative binding of EGF to an EGFR dimer. The peak value of 6 EGFR molecules per focus before EGF activation that we observe cannot be explained by a model as proposed previously^36^ which suggested that face-to-face dimers associate with the EGFR dimer interface between back-to-back dimers to generate higher order oligomeric complexes; analysis of the steady state solution for this model predicts a most likely stoichiometry of 4 EGFR per focus. Instead, a more likely state of 6 molecules (and higher after activation) suggests a more complex mechanism in which additional EGFR molecules result in greater stability for the overall cluster. This begs a question of what is the driving force behind cluster formation, which we do not directly address here. However, there is evidence from other studies that forces associated with molecular crowding in the membrane may result in oligomerization of integrated membrane proteins and the appearance of complex cytoskeletal and clathrin pit morphologies^34,55,62–64^. Ionic protein-lipid interactions^65^ and direct protein-protein interactions^33^ have also been implicated as contributory factors towards EGFR cluster generation.

Earlier work on heterocomplex formation in the Erb receptor family has suggested that EGFR and HER2 associate^31,34^, however, there are discrepancies in the interpretations of experimental data as to whether this association is before or after EGF activation. Our observations suggest that heterocomplex formation is most likely following EGF activation of EGFR. The physiological role of heterocomplex formation is unclear. HER2 is known to act as coreceptor but has no known direct ligand. However, upon transactivation (i.e. following activation of EGFR by EGF) it exhibits the highest of all kinase activities across the ErbB family^66^, thereby augmenting signalling efficiency. The mobility of heterocomplexes may potentially enable a spread of the signal across the surface of the cell, especially if HER2 molecules were to turn over between different EGFR complexes, however, this hypothesis remains to be tested. One consequence for having HER association post EGF binding is that the signal response at the level of the whole cell is more likely to be distinctly binary (i.e. highly biphasic) since the augmentation of the response due to HER2 association after activation results in a very high and rapid signal response. Our findings of post activation heterocomplex formation may suggest potential new strategies for anti-cancer drug design. For example, although there are anti-cancer drugs already established which bind specifically to HER2, one new strategy could be to target the specific interaction interfaces between HER2 and EGFR directly. Alternatively, it may also be valuable to explore new strategies to disrupt the oligomeric nature of the EGFR receptors before EGF activation.

## Methods

### Strain construction

We screened all ~100 colorectal cancer cell lines from the Cancer and Immunogenetics Laboratory (Weatherall Institute of Molecular Medicine, Oxford University, U.K.) for EGFR mRNA using available microarray data^42^ (Supplementary Fig. 1) and selected three preliminary lines (SW620, COLO320HSR and COLO741) on the basis of negligible native EGFR expression levels prior focussing on SW620 (COLO741 was found to be a melanoma line and COLO320HSR exhibited transfection instability). Total protein levels were estimated from cell lysates prepared from pellets using a radio immunoprecipitation assay lysis buffer supplemented with Roche cOmplete Mini ethylenediaminetetraacetic acid free protease inhibitor cocktail and Roche PhosSTOP phosphatase inhibitor cocktail. Total protein concentration was estimated using Thermo Scientific™ Pierce™ bicinchoninic acid Protein Assay Kit referenced against known concentrations of BSA. EGFR protein quantification was performed with western blotting, including cell lines with intermediate levels of EGFR expression as positive controls, probing nitrocellulose membranes with anti-EGFR mouse monoclonal antibody (1:1000, clone 1F4, Cell Signalling Technology^®^) and anti-β-tubulin mouse monoclonal antibody (1:1000, Sigma-Aldrich^®^) prepared in TBS-T, 5% milk and incubated overnight at 4ºC. After the washing, membranes were incubated with secondary antibody (1h, room temperature) using a polyclonal Rabbit anti-Mouse antibody conjugated to horseradish peroxidase (Dako) diluted at 1:10,000 and 1:100,000 for respectively EGFR and β-tubulin detection prior to Amersham Biosciences enhanced chemiluminescence (ECL) exposure.

Plasmid perbB1-EGFP-N1 (donated by Philippe Bastiaens, Max Planck Institute of Molecular Physiology, Dortmund, Germany) was used for transformations, which comprised an insertion of the human *EGFR* gene into the enhanced GFP Clontech backbone, pEGFPN1, plus selectable kanamycin *(kan)* resistance markers for bacterial/eukaryotic vectors. Competent *E. coli* cells were transformed with pEGFR-EGFP following the Invitrogen manufacturer’s protocol and plasmid DNA purified using the QIAGEN Plasmid Mini Kit. The concentration of purified DNA was determined using a Thermo Scientific NanoDrop™ 1000 Spectrophotometer at 230nm wavelength. SW620 cells were transfected with pEGFR-EGFP using Invitrogen’s cationic lipid transfection formulation, Lipofectamine^®^ LTX and Plus™ reagent. 1 day pre transfection 200,000 cells in 1ml growth medium were seeded into each well of a 12-well plate; the following day DNA-lipid complexes were prepared according to the manufacturer’s instructions. For each well we added 2μg plasmid DNA, 200μl Invitrogen Opti-MEM^®^ I Reduced Serum Medium, 1μl Plus Reagent and 6μl of Lipofectamine LTX. DNA-lipid complexes were added dropwise to the cells then placed in a 5% CO2 37 ºC incubator and the media changed after 5h to the usual cell media. The following day cells were trypsinized by trypsinization and reseeded onto 15cm plates in their usual growth media Gibco^®^ Dulbecco’s modified eagle medium (DMEM) supplemented with 4.5g/l glucose, pyruvate, L-glutamine and phenol red plus 2μg/ml Gibco™ Geneticin^®^ (G418 Sulphate) selection antibiotic. Once colonies were visible by naked eye, these were isolated using a silicon Cloning cylinder (Corning^®^), harvested using trypsin and transferred in a 12-well plate. Transgene expression was confirmed by three different methods of imaging live cells directly with confocal microscopy, imaging immunofluorescently stained fixed cells with confocal microscopy, and western blotting.

### Nanobody preparation

IgG antibodies to EGF and anti-EGF rabbit anti-mouse polyclonal IgG (Molecular Probes) were digested by papain, confirmed by migration of 28-30kDa and 25kDa molecular weight proteins under reducing conditions, corresponding to reduced Fc and Fab, respectively. Fab nanobodies were purified from the digest using protein A immobilized within a spin column. The completeness of IgG digestion and Fab purification were evaluated by measuring absorbance at 280nm wavelength using a Thermo Scientific NanoDrop spectrophotometer. Following purification of the digest a protein band at 25kDa only was seen in the protein A flow-through under denaturing and reducing conditions for both antibodies, consistent with reduced Fab.

### Fluorescence microscopy

For confocal microscopy we used a Zeiss inverted Axio Observer Z1 microscope with LSM 510 META scanning module and Plan-Aprochromat 63x 1.40NA oil immersion DIC M27 objective lens, enabling simultaneous imaging of green and red colour channels: excitation path used 488nm wavelength argon ion laser; first detection channel contained a 565nm wavelength dichroic beamsplitter and 505nm longpass emission filter for GFP, second channel collected transmitted light to produce a DIC image. Cells were grown in Corning 75 cm^2^ treated plastic cell culture flasks in a humidified incubator at 37 ºC with 5% carbon dioxide. Once cells were 70-100% confluent they were subcultured by enzymatic disaggregation with trypsin. 2-7 days prior to imaging, 150,000-300,000 SW620-EGFR-GFP cells were seeded onto Ibidi μ-dish 35mm, high glass bottom using their normal culture medium. SW620-EGFR-GFP cells were either seeded in DMEM containing phenol red, then changed to DMEM with addition of 4.5g/l glucose, L-glutamine, HEPES, without phenol red, and supplemented with 10% FBS, 100 units/ml of penicillin and 100 μg/ml of streptomycin, or SW620:EGFR-GFP cells were seeded directly into DMEM without phenol red. Prior to imaging the media was changed to Molecular Probes^®^ Live Cell Imaging Solution. All media used for SW620-EGFR-GFP cells were supplemented with 1.5mg/ml of G418 sulfate.

For immunofluorescent characterization we harvested SW620-EGFR-GFP cells 48h prior to fixation at density of ~50,000 per well seeded into Ibidi μ-Slide VI0.4, cultured in DMEM without phenol red, supplemented with 4.5g/l glucose, L-glutamine, HEPES 10% FBS and 100 units/ml of penicillin and 100μg/ml streptomycin, 1.5mg/ml of G418. Cells were fixed with 4% formaldehyde at room temperature for 10min and washed. Non-specific antibody adsorption was blocked with 10% FBS in PBS for 10-20min. Primary antibodies were EGFR (D38B1) XP rabbit monoclonal 4267P (Cell Signaling Technology, 1:50 dilution) and anti-GFP chicken IgY (H+L) (Cell Signaling Technology, 1:400 dilution) diluted in PBS with 10% FBS and 0.1% saponin overnight at 4 ºC. Each well was washed with 10% FBS and incubated with secondary antibodies, DyLight 633 goat anti-rabbit immunoglobulin G (IgG) highly cross adsorbed (PN35563, Thermoscientific), dilution 1:200, and Alexa Fluor 633 goat anti-chicken IgG (H+L) 2 mg/ml (Invitrogen) diluted in PBS with 10% FBS and 0.1% saponin. Channels were washed with PBS and Sigma Aldrich Mowiol 4-88 was added to solidify overnight. GFP, DyLight 633 or Alexa Fluor 633 and 4’,6-diamidino-2-phenylindole (DAPI) were individually illuminated and scanned. Transmitted light images were scanned simultaneously with GFP. GFP was excited as for live cell imaging, while DyLight 633 and Alexa Fluor 633 were excited by a 633nm HeNe laser and detection beam path contained a 565nm secondary dichroic beamsplitter and 650nm longpass filter.

For TIRF microscopy, a bespoke dual-colour single-molecule microscope was modified from a previous design^47,67,68^ equipped with nanostage (Mad City Labs), samples imaged at 37 ºC in a humidified stage top incubator supplemented with 5% CO_2_ (INUB-LPS, Tokai Hit). We used excitation sources of an Elforlight B4-40 473nm 40mW diode laser and Oxxius SLIM 561nm 200mW diode-pumped solid-state laser, independently attenutated and recombined into a common optical path prior to polarization circularization using an achromatic λ/4 plate before entering a Nikon Eclipse-Ti inverted microscope body. An achromatic doublet lens mounted onto a translation stage controlled the angle of incidence into the objective lens to generate TIRF via a Semrock 488/561nm BrightLine^®^ dual-edge laser-flat dichroic beam splitter into a Nikon CFI Apo TIRF 100x NA1.49 oil immersion objective lens. Our imaging system enabled simultaneous GFP and TMR detection across a laser excitation field of full width at half maximum 20μm laterally, intensity 1kW/cm^2^ and set depth of penetration ~100nm. Continuous fluorescence emissions were sampled at 30ms per frame and split into green/red channels via a 488/561nm dual-pass dichroic mirror (Semrock) and imaged onto two 512x512 pixel array EMCCD cameras (Andor, iXon+ DU-897 and iXon DU-887 for green and red respectively, piezoelectrically cooled to −70ºC) at ~50nm/pixel magnification, via Semrock 561nm StopLine^®^ single notch and Chroma 473nm notch filters. Cells were seeded and grown in culture medium onto glass-bottomed Petri dishes at 37 ºC in humidified 5% CO2, prior to imaging exchanging to Molecular Probes^®^ Live Cell Imaging Solution supplemented with G418 sulfate. When appropriate, EGF-TMR was added to stimulate activation, in addition to EGF inhibitors of cetuximab or trastuzumab (Molecular Probes) at a final concentration of 100 ng/ml. SW620:EGFR-GFP, and native SW620 cells with negligible endogenous EGFR as negative control, were imaged on plasma cleaned glass coverslips (25mm×75mm No. 1.5 D263M Schott) covered by a sterile Ibidi sticky-Slide VI0.4 in a laminar flow hood. 48h prior to imaging cells at a density of ~100,000 in a 50μl volume of DMEM without phenol red supplemented with 10% FBS, 100 units/ml Penicillin and 100μg/ml Streptomycin, were seeded into each channel of the slide. Prior to imaging the media was changed to DMEM without phenol red supplemented with 100 units/ml Penicillin and 100μg/ml Streptomycin but without the addition of FBS, supplemented with 1.5mg/ml G418 sulfate, serum starving the cells for ~12-24h prior to imaging to remove serum EGF. Although we cannot entirely exclude residual amounts of non-EGF EGFR ligands we checked the SW620 cell line for secretion of the most common ligands, indicating: EGF: not expressed; TGFA: low level expression; HBEGF: low level expression; AREG: not expressed; BTC: not expressed; EREG: not expressed; EPGN: no data available. Fluorescence image sequences and a brightfield image were acquired immediately after adding EGF-TMR (Molecular Probes) where appropriate to a final concentration of 100ng/ml, acquiring images at 5min intervals up to 60min.

For single-molecule *in vitro* TIRF we used surface-immobilized GFP or EGF-TMR via anti-GFP or anti-EGF antibodies (Molecular Probes) respectively followed by BSA to passivate the surface prior to washing off^69^. Whole IgG has in principle two binding sites and to test if two fluorophores may be seen in the same diffraction-limited fluorescent spot we also prepared an antigen binding fragment (Fab) with only one binding site. In brief, slides were constructed from Ibidi sticky-Slides VI0.4 and 25mm×75mm No. 1.5 D263M Schott glass coverslip. The coverslip was plasma-cleaned prior the 50μl of whole IgG or Fab applied to a single channel and incubated at room temperature for 5min. Channel then washed three times with 120μl of PBS and the remaining surface blocked with 50μl 1mg/ml of BSA for 60min. The channel was again washed three times with 120μl of PBS and then incubated with 50μl GFP for 7.5min or EGF-TMR for 4min. Finally the channel was washed five times with 120μl of PBS before applying 50μl 1:10000, 200nm diameter, 4% w/v, Invitrogen Molecular Probes carboxyl latex beads. These beads could be visualised in brightfield illumination for focussing to avoid using the GFP or TMR itself to focus on which would result in photobleaching. The slides were left 1-12h to allow latex beads to settle. Automated detection of fluorescent foci indicated no significant difference between brightnesses (Supplementary Fig. 3b) for the whole IgG or nanobody conjugation methods. We estimated the mean Gaussian sigma width for single GFP fluorescent foci images to be 230nm for GFP, a value which we interpret as the point spread function width of our imaging system corresponding to a peak emission wavelength of ~500nm.

### Foci tracking

Bespoke code written in MATLAB (Mathworks)^44,47^ was used to track single fluorescent foci in green and red channels to determine spatial localization and calculate integrated foci pixel intensities and microscopic diffusion coefficients. The centroid of each fluorescent focus is determined using iterative Gaussian masking to a sub-pixel precision of ~40nm. The focus brightness is calculated as the sum of the pixel intensities inside a 5-pixel-radius region centred on the centroid, after subtraction of local background intensity. The signal-to-noise ratio (SNR) for a fluorescent focus is defined as the total focus intensity per pixel divided by the standard deviation of the background intensity per pixel. When the SNR for a focus is >0.3, the focus is accepted and fitted with a 2D radial Gaussian function to determine its Gaussian sigma width. We decided on an SNR threshold level of 0.3 as a compromise between a high probability for true positive detection but a low likelihood for false positive detection at single-molecule fluorophore intensity levels. We simulated fluorescent foci as 2D Gaussian functions in bespoke code written in MATLAB with comparable integrated pixel intensity values and widths as for those measured experimentally for single GFP/TMR used in the surface-immobilization assays, adding similar levels of Poisson-distributed noise, and ran these synthetic data through the same foci detection algorithms as for real experimental data, but exploring a range of SNR detection thresholds. We found that a threshold of 0.3 gave a true position detection probability of approaching 50% over a signal range corresponding to 1-10 fluorophores per focus, but with a false positive detection probability an order of magnitude less.

Foci detected in the tracking algorithm in consecutive image frames separated by 5 pixels or less (approximately the point spread function width of our imaging system), and which are not different in brightness or sigma width by more than a factor of two, are linked into the same track.

### Stoichiometry analysis

Stoichiometry per fluorescent focus was estimated in bespoke code written in MATLAB using integrated intensities and step-wise fluorophore photobleaching with Fourier spectral analysis to determine the brightness of either GFP or TMR during live cell imaging^69^. The brightness of a single GFP or TMR in our microscope was determined from *in vivo* data and corroborated using *in vitro* immobilised protein assays. The brightness of tracked foci in live cells followed an approximately exponential photobleach decay function of intensity with respect to time. Every fluorescent foci as it photobleaches to zero intensity will emit the characteristic single GFP brightness value, *I_GFP_*, in the case of EGFR-GFP, and *I_TMR_* in the case of EGF-TMR, given in our case by the modal value of all spot intensities over time, and can bleach in integer steps of this value at each sampling time point. Estimates for *I_GFP_* and *I_TMR_* were further verified by Fourier spectral analysis of the pairwise distance distribution^69^ of all spot intensities which yielded the same value to within measurement error. The initial intensity *I_0_* was estimated by interpolation of the first 3 measured data points in each focus track. Stoichiometries were obtained by dividing *I_0_* by of a given focus track by the appropriate single-molecule fluorophore brightness. Stoichiometry distributions were rendered as Gaussian kernel density estimations^69^ using standard MATLAB routines.

### Mobility analysis

For each accepted focus track, the 2D mean square displacement (MSD) was calculated in bespoke code written in MATLAB from the fitted focus centroid at time *t*, (*x*(*t*),y(*t*)), assuming a track of *N* consecutive image frames, and a time interval τ = *n*Δ*t*, where *n* is a positive integer and Δ*t* the frame integration time^70^:

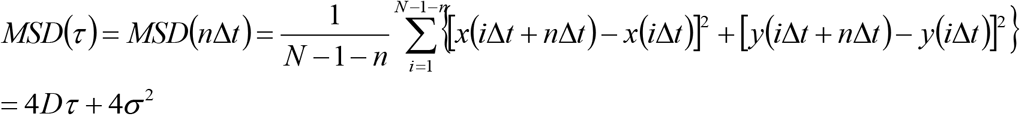

The localisation precision from our tracking algorithm (i.e. on the x-y image plane) is given by σ, which we estimate as 40 *±* 20nm. The apparent diffusion coefficient *D* is estimated from a linear fit to the first three data points in the MSD curve as a function of τ (i.e. 1 ≤ *n* ≤ 3, corresponding to the linear region of the average MSD *vs*. τ plot) for each accepted track, the fit constrained to pass through a point 4*σ*^2^ on the vertical axis corresponding to τ = 0, allowing *σ* to vary in the range 20 - 60nm in line with the measured range during the fitting optimisation.

### Colocalization analysis

The extent of colocalization was quantified using foci overlap integration between green and red channels^47^ determined by calculating the overlap integral between each green/red pair in bespoke code written in MATLAB, whose centroids were within 5 pixels of each other (i.e. green/red pairs in close proximity). In brief, assuming two normalized, 2D Gaussian intensity distributions *g*_1_(*x,y*) and *g*_2_(*x,y*), centered around (*x*_1_,*y*_1_) with width *σ*_1_, and centred around (*x*_2_,*y*_2_) with width *σ*_2_ for green and red foci respectively, the overlap integral *v* can be analytically calculated as^47^:

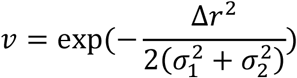

where:

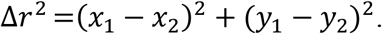

Our simulations indicate that a green/red foci pair that have identical centroid coordinates can have a measured overlap integral as low as ~0.75 due to the finite localization precision of 40nm. Therefore, we used an overlap integral threshold of ≥0.75 as a criterion for colocalization for the experimental data.

### Modelling the overlap probability of EGFR-GFP foci images

The probability that two or more fluorescent foci are within the diffraction limit of our microscope was determined in bespoke code written in MATLAB using a previously reported model^47^ at foci surface density values observed here. Such overlapping foci are detected as higher apparent stoichiometry foci. The stoichiometry distribution from overlapping foci was modelled by convolving a Poisson distribution generated from the probability of overlap with the expected intensity distribution of an isolated multimer. The latter is obtained by scaling the width of the single fluorophore intensity distribution (Supplementary Fig. 3) by *S*^1/2^ where *S* is the model stoichiometry. This model stoichiometry was fixed for those shown in Supplementary Fig. 5. For the Monte Carlo model, the model stoichiometry was generated from a population distribution of oligomeric EGFR whose stoichiometry was sampled from a random Poisson distribution with mean value equal to the mode peak value of 6 that we observed. This prediction resulted in a reasonable fit to the experimental distribution with goodness-of-fit *R*^2^=0.4923.

### Software access

All our bespoke code written in MATLAB is available from file EGFRanalyser at https://sourceforge.net/projects/york-biophysics/.

### Statistical tests and replicates

Two-tailed Student *t*-tests were performing for comparisons between pairs of datasets to test the null hypothesis that data in each was sampled from the same statistical distribution. We assume (n_1_+n_2_-2) degrees of freedom where n_1_ and n_2_ are the number of independent data points in each distribution and by convention that *t* statistic values which have a probability of confidence P>0.05 are statistically not significant. For TIRF each cell was defined as a biological replicate sampled from the cell population. We chose sample sizes of 10-117 cells per experimental condition which generated reasonable estimates for the stoichiometry distributions. Technical replicates are not possible with irreversible photobleaching, nevertheless, the noise in all light microscopy experiments has been independently characterized for the imaging system used previously.

## Acknowledgements

We thank Philippe Bastiaens, Max Planck Institute of Molecular Physiology, Dortmund, Germany for donation of plasmid perbB1-EGFP-N1. Work was supported by the EPSRC (EP/G061009/1), Royal Society (RG0803569, UF110111), BBSRC (BB/F021224/1, BB/N006453/1), MRC (MR/K01580X/1, PhD studentship) and CRUK (C38302/A12278).

---

**Supplementary Figure 1.**
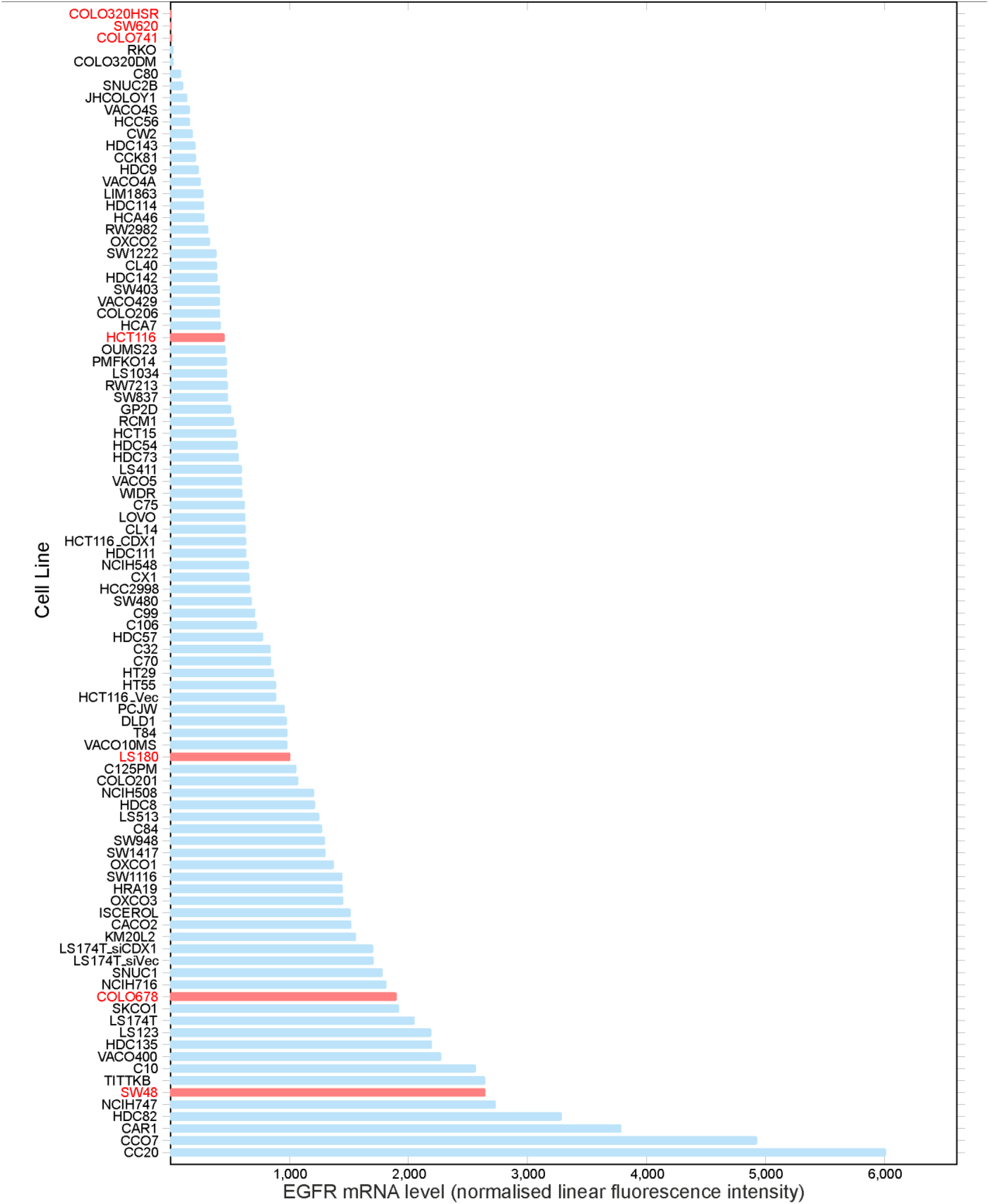
EGFR mRNA expression levels. Expression levels were quantified for a colorectal cancer cell line panel using Affymetrix U133+2 mRNA microarray data. Measurements indicated three candidate cell lines, SW620, COLO320HSR and COLO741 (labelled in red, top of panel), as having very low levels of native EGFR expression, as tested in subsequent western blot analysis in comparison to EGFR-expressing cell lines as positive controls (indicated as red columns, middle and bottom of panel). Three candidate cell lines with very low or absent levels of EGFR mRNA (SW620, COLO320HSR, COLO741; Y axis text label in red, top of panel) and a further four positive controls with medium to high levels (HCT116, LS180, COLO678; indicated as red columns, middle and bottom of panel), were selected and protein levels confirmed by western blot (Figure 1b).

**Supplementary Figure 2.**
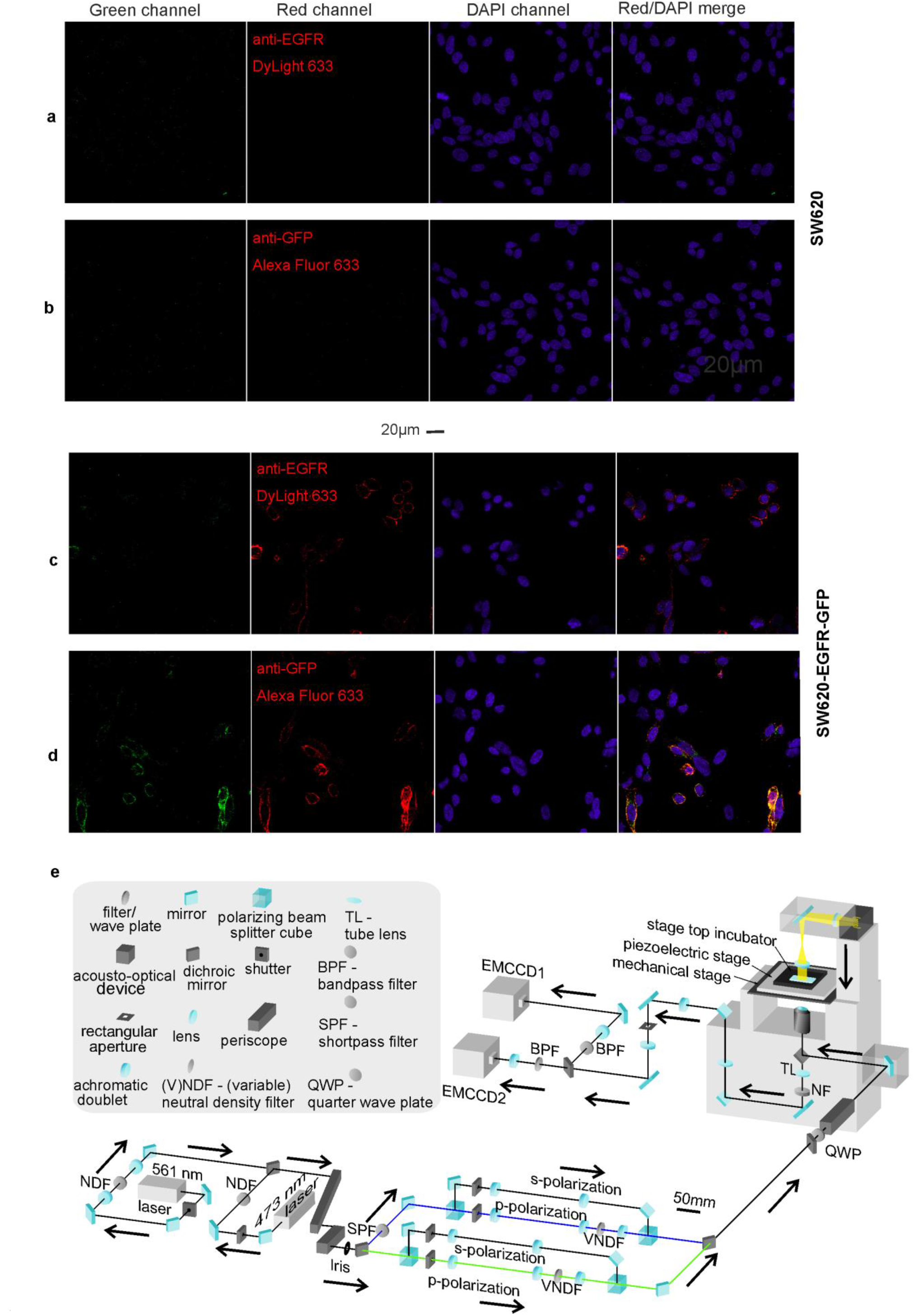
Confocal and TIRF characterization. Confocal microscopy images of fixed cells using GFP, anti-GFP immunofluorescence, and DAPI staining: (**a,b**) non-GFP background cell line SW620; (**c,d**) SW620-EGFR-GFP; (**e**) optical path diagram of bespoke single-molecule TIRF microscope.

**Supplementary Figure 3.**
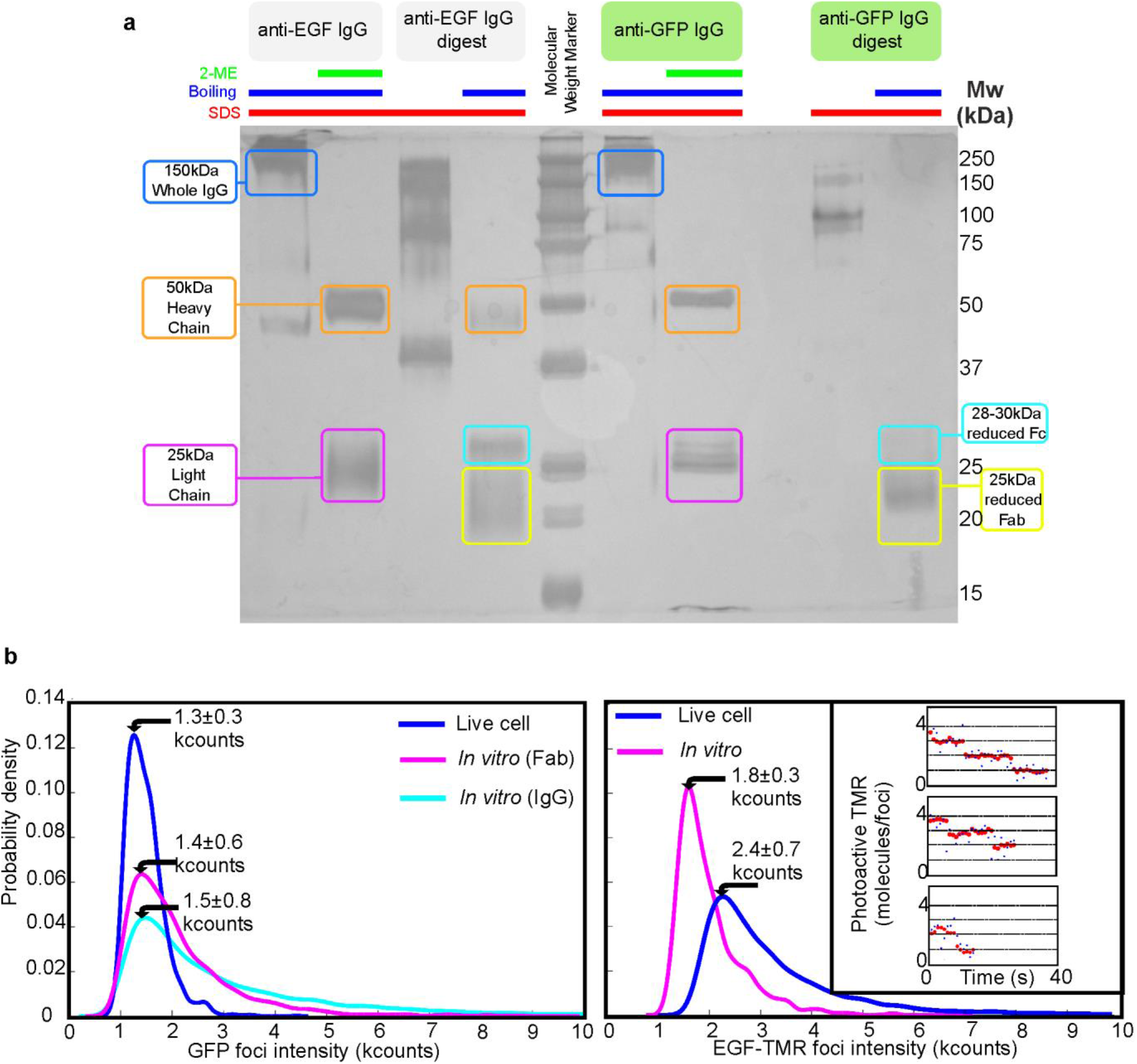
Characterization of unitary fluorophore brightness values. (**a**) SDS-PAGE gel indicating generation of Fab nanobody fragments (yellow) from anti-EGF and anti-GFP IgG antibodies (blue), heaving (orange) and light chains (magenta) indicated with reduced Fc (cyan). (**b**) Kernel density estimation^69^ distributions of fluorescent foci intensity values measured in kcounts (i.e. counts x 10^3^) for single GFP (left panel) for live cell, at the end of the photobleach, before EGF is added compared with *in vitro* Fab and whole IgG data. TMR molecule data for *in vitro* EGF-TMR and live cell, at the end of the photobleach, post EGF binding data taken from colocalized EGF-EGFR foci is shown (right panel); inset shows live cell EGF-TMR photobleach steps after EGF has been added, taken from colocalized EGF-EGFR foci, with raw (blue) and Chung-Kennedy filter^48,49^ (red) traces, mean and s.e.m. indicates (arrows).

**Supplementary Figure 4.**
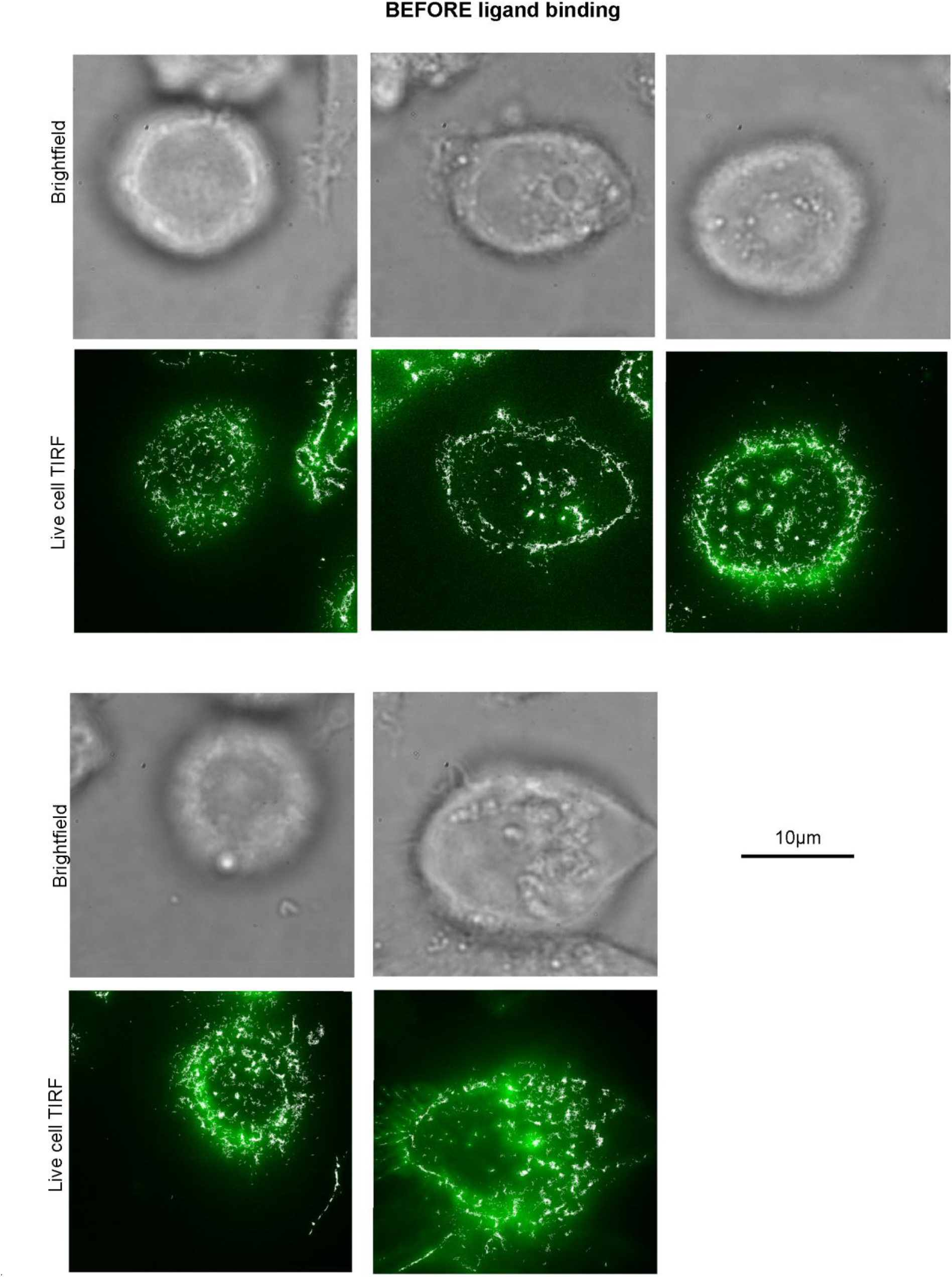
More examples of cells before addition of EGF ligand. Brightfield images (grey) and TIRF (green) shown with overlaid foci tracking output (white).

**Supplementary Figure 5.**
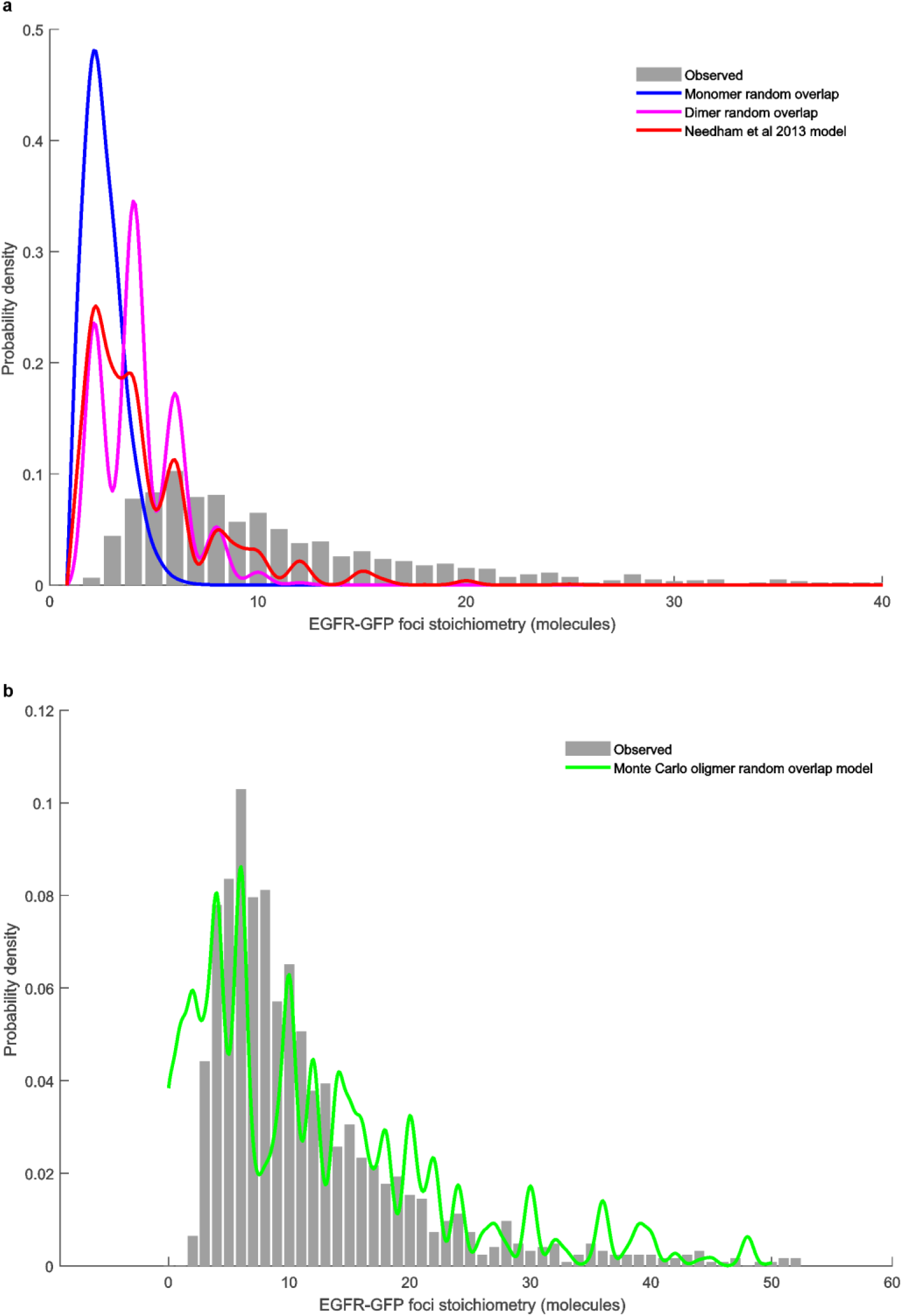
Random foci overlap model. Random overlapping foci predictions for (**a**) monomeric (blue) and dimeric EGFR (magenta), and a mixed model oligomer model suggested from a previous single-molecule study (red)^35^, all showing poor agreement (R^2^<0) to our experimental observations for stoichiometry distribution (grey). (**b**) Monte Carlo Poisson model using an expected average value of 6 molecules for EGFR foci stoichiometry (green) showing reasonable fit (R^2^=0.4923) to experimental data (grey).

**Supplementary Figure 6.**
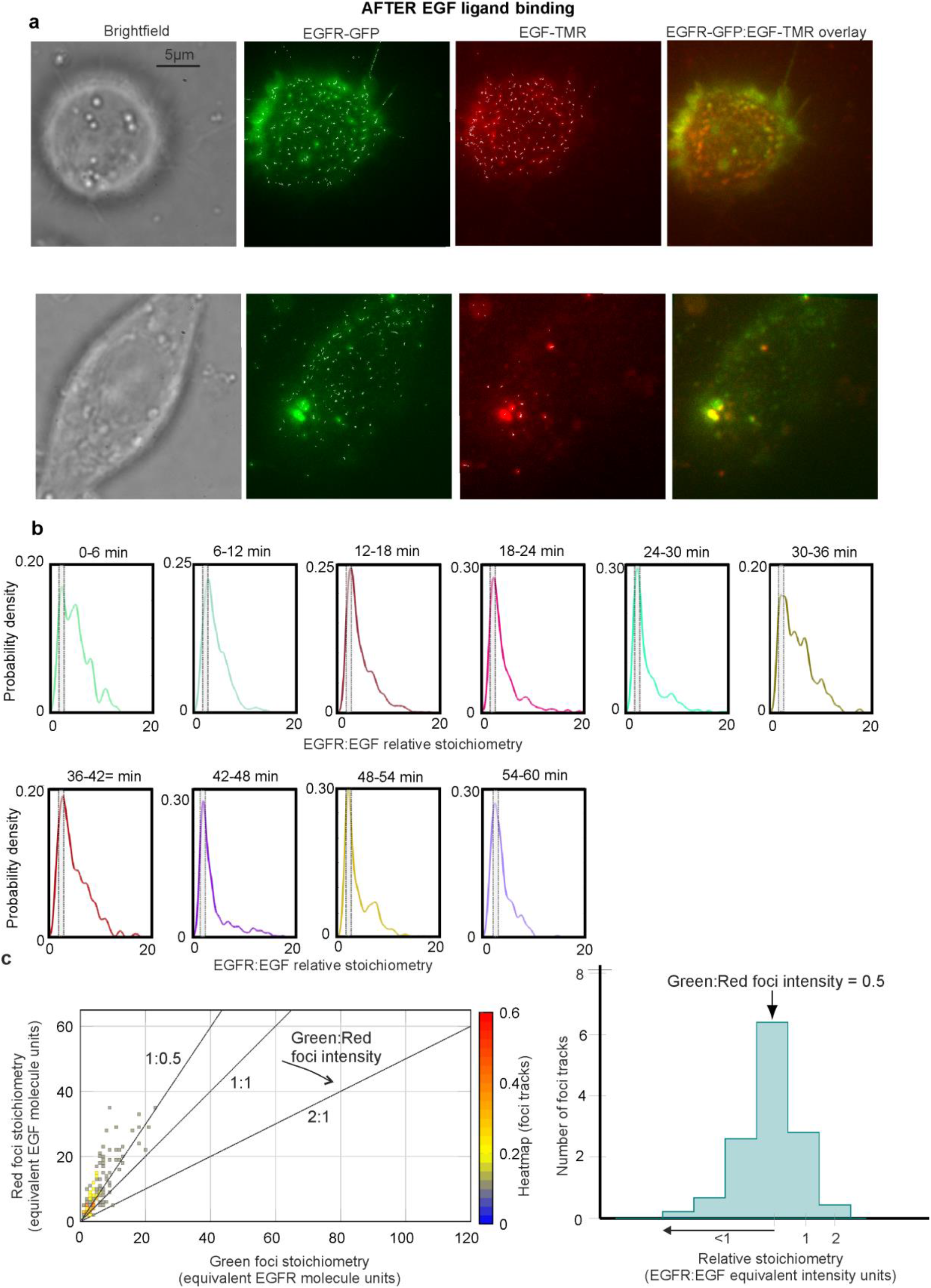
Charactering EGFR and EGF foci stoichiometry after addition of EGF. (**a**) Two examples of cells taken ~10 min after the addition of EGF: brightfield images (grey), green channel showing EGFR-GFP localization (green), red channel showing EGF-TMR localization (red), and the overlay of green and red channels together (right panels, with yellow indicating regions of high colocalization) are shown here. (**b**) Variation of the EGFR:EGF relative stoichiometry, rendered as kernel density estimations, as a function of incubation time with EGF (shown in 6 min bins). The region corresponding to 2.0 ± 0.5 relative stoichiometry is indicated as a grey rectangle. (**c**) Heatmap (left panel) and histogram (right panel) characterizing ‘false’ colocalization due to cellular autofluorescence in green and red channels.

**Supplementary Figure 7.**
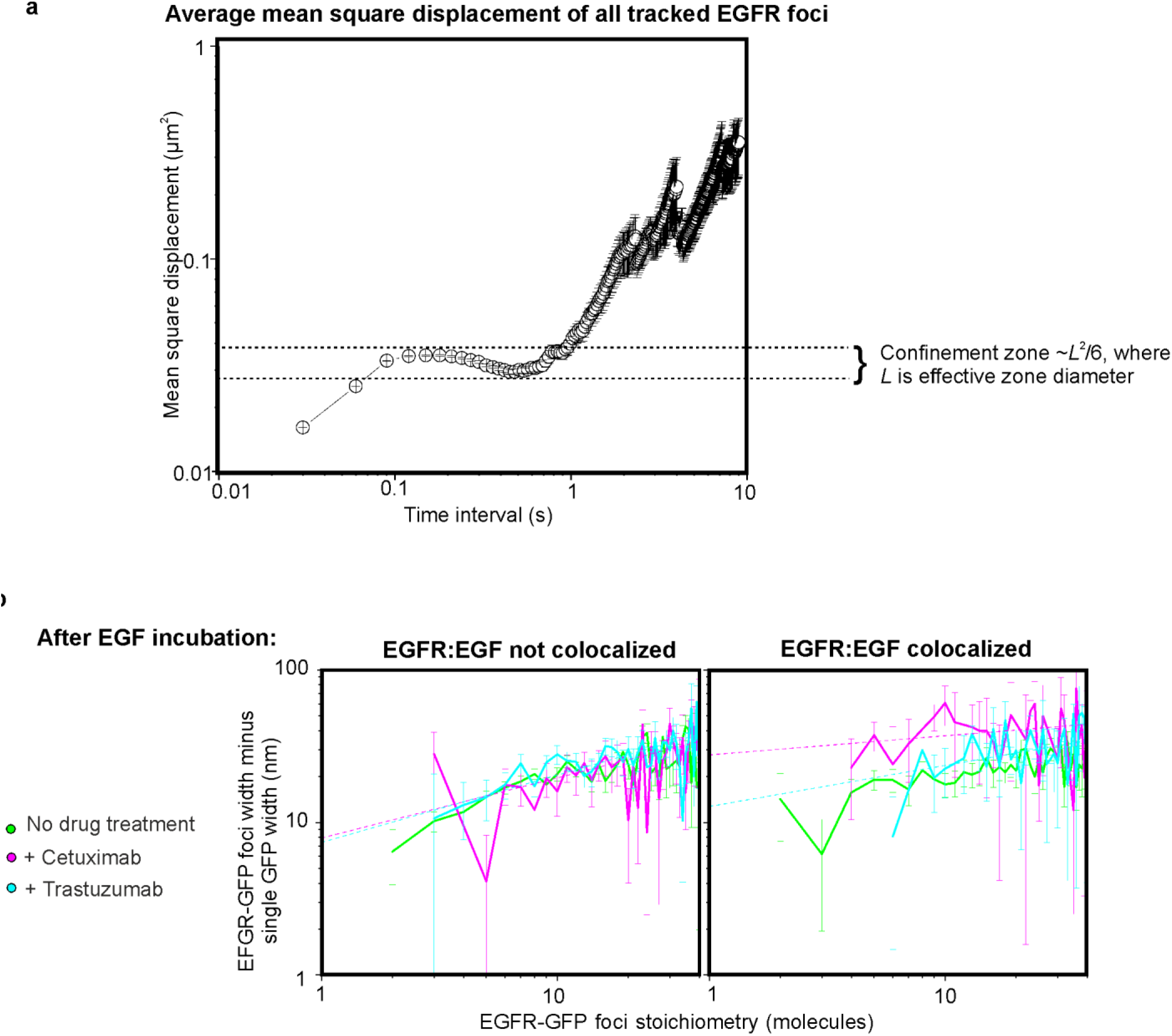
EGFR foci diffusion. (**a**) Log-log plot for average mean square displacement *vs*. time interval for all collated EGFR-GFP foci tracks before addition of EGF, putative confinement zone indicated (dashed lines), from number of foci N=770, acquired from number of cells N=19. (**b**) Log-log plots of EGFR-GFP foci diameters minus the width for a single GFP molecule *vs*. stoichiometry for not colocalized (left panel) and colocalized foci (right panel), showing cells with no cetuximab or trastuzumab treatment (green, N=6,710 foci, N=117 cells), those treated with cetuximab (magenta, N=1,219 foci, N=25 cells), and those treated with trastuzumab (cyan, N=1,607 foci, N=27 cells), with heuristic power law fit (dash lines), s.e.m. error bars.

